# SOX9 binds TCFs to mediate Wnt/β-catenin target gene activation

**DOI:** 10.1101/2022.04.06.487337

**Authors:** Aravinda-Bharathi Ramakrishnan, Peter E. Burby, Kavya Adiga, Ken M. Cadigan

**Affiliations:** Department of Molecular, Cellular, and Developmental Biology, University of Michigan, Ann Arbor, MI 48109 USA

## Abstract

Activation of the Wnt/β-catenin pathway regulates gene expression by promoting the formation of a β-catenin-TCF complex on target gene chromatin. The transcription factor SOX9 antagonises Wnt signalling in many contexts through its ability to down-regulate β-catenin protein levels. Here, we find that SOX9 promotes the proliferation of Wnt-driven colorectal cancer (CRC) cells. We identify multiple enhancers activated by SOX9 working in concert with the Wnt pathway. These enhancers are co-occupied by TCFs and SOX9 and contain TCF and SOX9 binding sites that are necessary for transcriptional activation. In addition, we identify a physical interaction between the DNA-binding domains of TCFs and SOX9 and show that TCF-SOX9 interactions are important for target gene regulation and CRC cell growth. Our work demonstrates a highly context-dependent effect of SOX9 on Wnt targets, with activation or repression being dependent on the presence or absence of SOX9 binding sites on Wnt-regulated enhancers respectively.

## Introduction

Transcriptional regulation by the Wnt/β-catenin pathway is essential for metazoan development and homeostasis (*1*), and the transcription factors (TFs) of the TCF/LEF family (TCFs) are major effectors of this signalling cascade (*2*). Clusters of TCF binding sites are characteristic of many WREs (Wnt-regulated elements), the *cis*-regulatory DNA elements that drive Wnt-responsive transcription (*3*). WRE activation is triggered by the Wnt-dependent stabilisation and accumulation of the protein β-catenin, which is recruited to WREs by TCFs. β-catenin recruits co-activators to induce Wnt target gene transcription (*2, 4*).

While TCFs mediate Wnt target gene activation across cell types, there is great spatio-temporal diversity in the identity of Wnt target genes, with different tissues expressing unique cell-type specific Wnt transcriptional programs. This specificity is thought to arise from interactions between TCFs and other TFs that help determine cell fate and identity. Many non-TCF TFs bind to WREs and affect Wnt target gene expression, but a comprehensive molecular understanding of how these TFs generate target gene specificity is still incomplete (*5, 6*). One well-studied example is TFs of the CDX family, which bind to several WREs and recruit TCFs to them (*7, 8*). Their ability to form complexes with TCFs is essential for the activation of several Wnt target genes (*9*).

Another group of developmentally important TFs that modulate Wnt target gene expression is the SOX family of proteins. SOX3, SOX4, and SOX17 directly bind to TCFs and β-catenin and promote the expression of specific Wnt targets (*10, 11*). In diverse contexts such as *Xenopus* endoderm differentiation and in human pluripotent stem cell (hPSC) differentiation, SOX17 and β-catenin co-occupy and co-regulate a variety of WREs. Analyses of chromatin occupancy suggest that for a subset of these WREs, SOX17 is required for β-catenin recruitment (*12, 13*).

SOX proteins are also known to repress Wnt signalling, with SOX9 being the most intensively studied. SOX9 has essential functions in the development of cartilage and the skeletal system, and mutations in SOX9 are associated with sex reversal and severe skeletal deformities in humans (*14, 15*). SOX9 is also well-known for its role in promoting testis formation during mammalian development (*16*). Genetic studies have shown that testis or ovary commitment in mammalian gonadal development involves a mutual antagonism between the Wnt pathway, which promotes ovarian development and inhibits testis fate, and SOX9, which inhibits the Wnt pathway to promote testes growth (*17*). In several different mammalian cell lines, the overexpression of SOX9 causes a reduction in Wnt transcriptional readouts and in β-catenin protein levels (*18*–*21*). Recent work from our group found that SOX9 promoted β-catenin degradation in a destruction complex and proteasome-independent manner by transcriptionally activating the Notch pathway coactivator MAML2, which associates with β-catenin (*20*).

Although primarily characterised as a Wnt antagonist, correlative evidence hints at a more complex relationship between SOX9 and Wnt signalling in the intestinal epithelium and in colorectal cancer (CRC). The villi of the intestinal epithelium consist of short-lived terminally differentiated cells that are continuously replenished from a population of intestinal epithelial stem cells. The proliferation of these stem cells and the differentiation of the Paneth cells which surround and metabolically support them are both dependent on Wnt signalling (*22, 23*). Paneth cells express high levels of SOX9 (*24*), and the loss of either SOX9 or Wnt signalling prevents Paneth cell differentiation (*22, 25, 26*). Additionally, the overexpression of SOX9 in Wnt-dependent CRC lines induces the expression of many genes characteristic of Paneth cells (*27*). Some of these are also directly activated by Wnt signalling (*27*–*29*), suggesting the possibility of SOX9 and Wnt signalling working together to activate gene expression. Understanding the relationship between SOX9 and Wnt signalling in this context requires an understanding of the regulatory logic of target gene WREs regulated by both factors.

Aberrant Wnt pathway activation in intestinal stem cells leads to uncontrolled proliferation, and consequently, activating mutations in the Wnt pathway are major drivers of CRC (*4, 30*). Additionally, mutations that increase the activity of WREs that regulate oncogenes such as *MYC* are also associated with increased cancer risk (*31*–*34*). Many Wnt-dependent CRC lines and primary CRC samples also show high levels of SOX9 expression (*35, 36*), and recent reports suggest that SOX9 promotes stemness and survival of some CRC lines (*27, 37, 38*). Findings of SOX9 promoting Paneth cell fate and CRC cell survival are at odds with models of SOX9 as a Wnt antagonist.

In this study, we interrogate the relationship between SOX9 and Wnt signalling in regulating gene expression. We show that SOX9 and Wnt signalling work together to promote the growth and survival of CRC cells. Among the target genes activated by Wnt signalling and SOX9 is the oncogene *MYC*. In CRC cells, SOX9 is associated with many WREs, including several cancer risk-associated enhancers of *MYC*. Our characterisation of the regulatory logic c-Myc-335 WRE reveals the presence of SOX9 binding sites, which regulate enhancer activity along with TCF sites. A similar regulatory logic is also seen in the promoters of *Defa5* and *Defa6*, two markers of Paneth cells, that are synergistically upregulated under high Wnt, high SOX9 conditions. With reporter mutagenesis and synthetic reporters, we show that the combination of TCF and SOX binding sites is necessary and sufficient for synergistic activation by Wnt and SOX9. Mechanistically, we show that SOX9 directly binds to TCFs. Through a novel separation-of-function mutant, we show that the activation of WREs and CRC cell growth both require not just the activities of TCFs and SOX9, but also the activity of a TCF-SOX9 complex. Our work demonstrates that in addition to its role as a Wnt antagonist, SOX9 works together with the Wnt/β-catenin pathway to activate a subset of Wnt target genes.

## Results

### SOX9 and Wnt signalling cooperate to promote the growth and survival of colorectal cancer cells

To explore the roles of SOX9 and Wnt signalling in CRC, we tested the consequences of depleting SOX9 and/or β-catenin in LS174T CRC cells. These cells contain activating mutations in β-catenin and are dependent on Wnt signalling for their growth and survival (*39*). We used a previously engineered LS174T cells containing a doxycycline (DOX)-inducible β-catenin shRNA expression cassette (*39*) (referred to as LS174T-pTER-β-cat cells), and transduced them with constructs expressing either non-targeting (Scrambled) or SOX9 targeting shRNAs (Table S1). Western blots showed reduced SOX9 expression in the two SOX9 shRNA lines (Fig. 1A, lanes 2,3). This reduction of SOX9 protein levels was not accompanied by an upregulation of β-catenin protein, as would be expected if SOX9 was promoting β-catenin turnover (Fig. 1A, lanes 2,3). This suggested that SOX9 is not a β-catenin antagonist in these cells. SOX9 protein levels were not affected by β-catenin depletion, suggesting that SOX9 is not a target of Wnt signalling in these cells (Fig. 1A, lanes 1,4).

**Figure 1.**
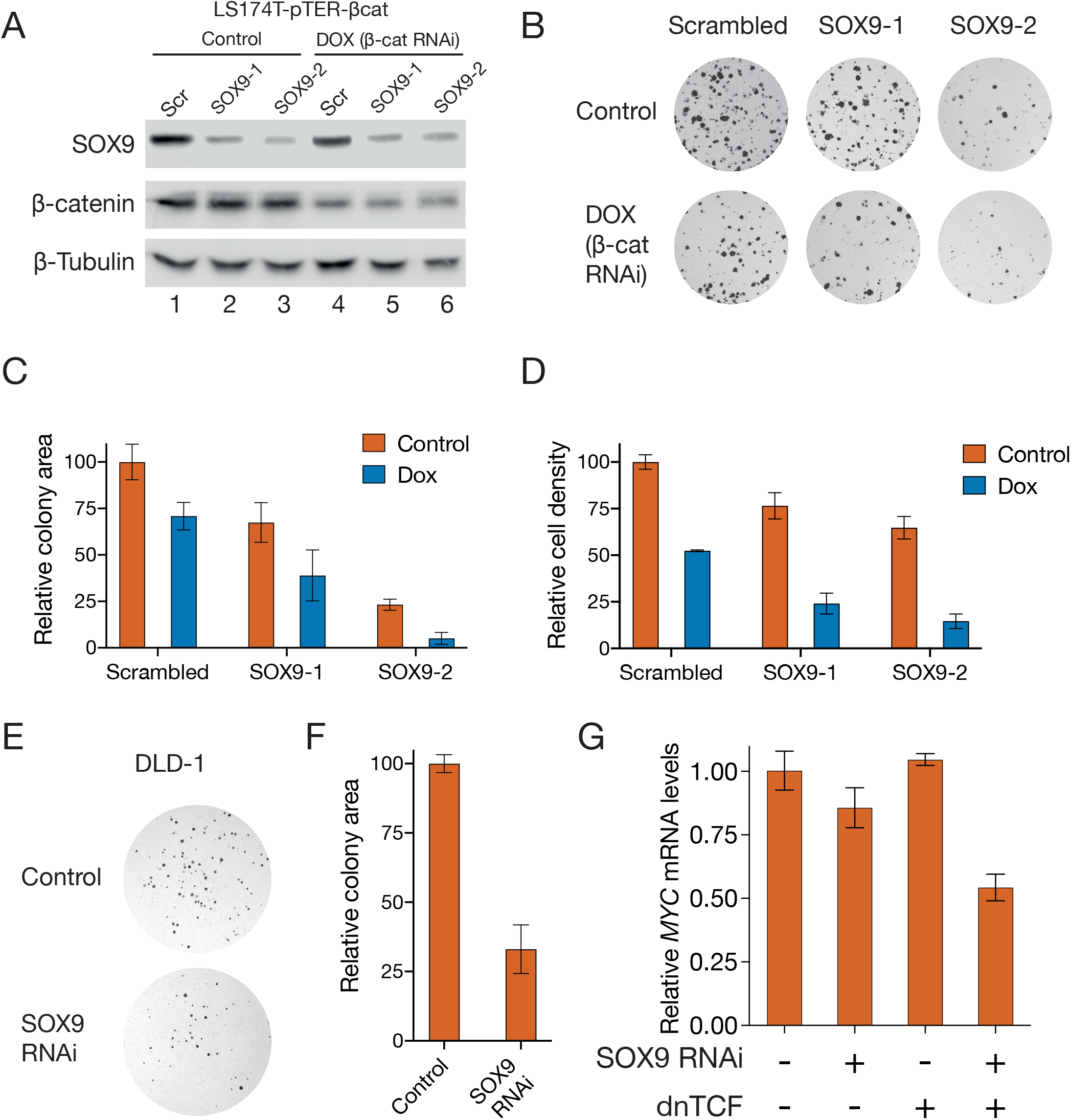
Wnt signalling and SOX9 work together to promote the growth and proliferation of colorectal cancer cells. **A)** Western blots showing depletion of SOX9 and β-catenin in LS174T colorectal cancer cells by RNAi. Cells were stably transduced with constructs expressing either a scrambled (Scr) or two independent SOX9-targeting shRNAs (SOX9-1, SOX9-2) and then treated with DOX for 24h to deplete β-catenin. **B)** Representative images showing reduced colony formation in LS174T cells after knockdown of SOX9 and β-catenin. Cells were treated in similar conditions as in (A) and then seeded at a density of 1000 cells/well. **C)** Quantification of the area occupied by the colonies shown in (B). **D)** RNAi against β-catenin and SOX9 reduces the growth rate of LS174T cells. Cells were treated with DOX for 48h, following which relative cell numbers were measured using an MTT assay. **E)** Reduced colony formation in DLD-1 colorectal cancer cells after SOX9 knockdown. Cells were transfected with either a control plasmid or one expressing a SOX9 targeting shRNA and plated at a density of 500 cells/well. **F)** Quantification of the area occupied by the colonies shown in (E). **G)** RT-qPCR data showing *MYC* mRNA levels in DLD-1 cells after transfection with plasmids expressing dominant negative TCF (dnTCF), SOX9 RNAi, or both. mRNA levels were measured with RT-qPCR 24h after transfection.

To understand whether SOX9 and Wnt signalling worked together to promote the survival of LS174T-pTER-β-cat cells, we tested for a SOX9 requirement in a colony formation assay with and without DOX-induced β-catenin depletion. Knocking down SOX9 alone resulted in fewer colonies, and the combined depletion of β-catenin and SOX9 reduced the clonogenicity of these cells even further (Fig. 1B,C). We then tested the impact of Wnt signalling and SOX9 on the growth rate of these cells using an MTT assay. The depletion of β-catenin or SOX9 led to slower growth, and combined depletion of β-catenin and SOX9 reduced growth four-to-five fold (Fig. 1D). To ensure that our findings were not specific to LS174T cells, we also tested the role of SOX9 in promoting the survival of DLD-1 cells. DLD-1 cells also have elevated levels of Wnt signalling, but unlike LS174T cells, express a WT β-catenin protein and a truncated form of the destruction complex component APC (*40*). In DLD-1 cells, SOX9 depletion caused a three-fold reduction in colony formation (Fig. 1E,F). Just like in LS174T cells, SOX9 depletion did not cause an upregulation of β-catenin protein levels, and the inhibition of Wnt signalling through a dominant negative TCF (dnTCF) construct (*41*) did not reduce SOX9 protein levels (Fig. S1A).

We then examined the importance of Wnt signalling and SOX9 across a panel of CRC lines using data from the Cancer Cell Line Encyclopedia (CCLE) (*42, 43*). First, we examined gene expression data and correlated it with dependency scores calculated from CRISPR knock-out experiments using the Chronos algorithm (*44*). As expected in CRC, 49/53 lines examined showed high β-catenin (*CTNNB1*) expression and dependence based on our cutoffs (Fig. S1B). A similar analysis for SOX9 revealed that 38/53 lines showed high SOX9 expression and SOX9-dependent growth (Fig. S1C). Dependence on SOX9 was highly correlated with β-catenin-dependence, with 40/56 lines being highly dependent on both β-catenin and SOX9 (Fig. S1D). As a negative control, we looked at SOX17, which represses Wnt targets and inhibits the proliferation of CRC cells (*11*). Just 2/56 lines showed a dependence on both genes (Fig. S1E). Across all cancers in the CCLE database (n=1070), β-catenin was the seventh-most correlated gene with SOX9 in terms of essentiality (Fig. S1F). In conjunction with our experimental data, these analyses strongly indicated that Wnt signalling and SOX9 working together was a general feature of CRC.

Since both Wnt signalling and SOX9 are major regulators of transcription, we wondered whether SOX9 could cooperate with the Wnt pathway to activate a common transcriptional program in CRC. One of the best-characterised Wnt targets in the context of CRC is the *MYC* oncogene (*45*), which is regulated by several WREs associated with CRC risk (*31*). In DLD-1 cells, *MYC* transcripts were downregulated by the combination of SOX9 knockdown and Wnt inhibition by dnTCF (Fig. 1G).

### SOX9 directly binds and regulates the CRC risk-associated c-Myc-335 WRE

After finding that *MYC* transcript levels were upregulated by Wnt signalling and SOX9, we wanted to study the regulatory interactions that allowed SOX9 to activate this Wnt target. One of the best-characterised WREs regulating *MYC* is the c-Myc-335 enhancer containing a SNP (rs6983267) that increases CRC risk in humans (*9, 46, 47*). Examination of a previously published SOX9 ChIP-seq dataset (*48*), showed that c-Myc-335 was bound by SOX9 in HT-29 CRC cells (Fig. 2A). We verified this result in LS174T cells by performing ChIP using antibodies against TCF7L2, β-catenin, and SOX9. To detect ChIP enrichment, we compared qPCR signals from primers located inside the enhancer to those from two primer sets flanking the enhancer (Fig. 2B), and found that TCF7L2, β-catenin, and SOX9 were all enriched at the enhancer (Fig. 2C).

**Figure 2.**
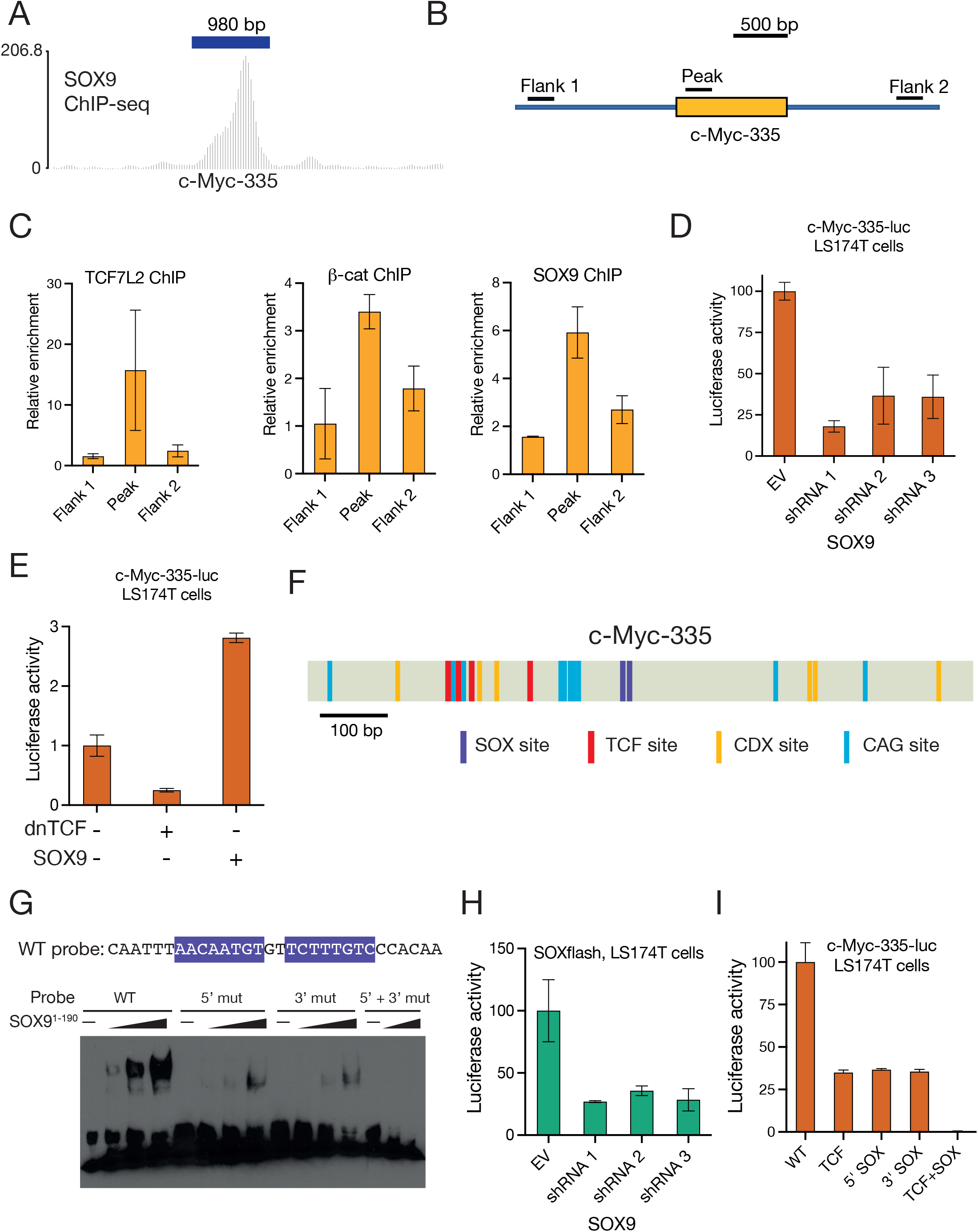
The colorectal cancer-risk associated c-Myc-335 WRE requires the direct binding of TCFs and SOX9 for its activation. **A)** ChIP-seq tracks showing the binding of SOX9 to the c-Myc-335 locus in HT29 colorectal cancer cells. Dataset from (*48*). **B)** Cartoon showing positions of primers used for ChIP-qPCR analysis of the c-Myc-335 locus in LS174T colorectal cancer cells. **C)** ChIP-qPCR showing TCF7L2, β-catenin, and SOX9 are enriched at the c-Myc-335 locus in LS174T cells. ChIP signals were quantified by qPCR using primers shown in (B). **D)** RNAi against SOX9 reduces c-Myc-335 reporter activity in LS174T cells. Cells were transfected with the reporter plasmid and plasmids encoding shRNAs against SOX9 or an empty vector (pSUPER). Luciferase activity was assayed 48h post transfection. **E)** Luciferase assay showing the effect of dnTCF and SOX9 overexpression on c-Myc-335 reporter activity in LS174T cells. Cells were transfected with the reporter and plasmids encoding dnTCF, Flag-SOX9, or an empty vector (pcDNA3.1). Luciferase activity was assayed 24h post transfection. **F)** Cartoon of the c-Myc-335 enhancer showing positions of the newly identified SOX binding sites (purple) along with previously characterised TCF, CDX, and CAG sites (*9*). **G)** EMSA showing SOX9 specifically binds to the newly-identified binding sites on c-Myc-335, and mutations in these sites abolish SOX9 binding. **H)** Luciferase assay showing the effect of SOX9 RNAi on the activity of a synthetic reporter containing 3 tandem copies of the SOX9 binding sites from c-Myc-335. Reporter was measured similarly as in (D). **I)** Luciferase assay showing the effect of TCF and SOX site mutations on c-Myc-335 reporter activity. WT and mutant reporters were transfected into LS174T cells and luciferase activity was assayed 24h post transfection.

Previous work from our group had generated a luciferase reporter driven by c-Myc-335 which showed Wnt-dependent activity (*9*). In LS174T cells, we found that RNAi-mediated SOX9 depletion reduced c-Myc-335 reporter activity (Fig. 2D). Wnt pathway inhibition with dnTCF repressed, while SOX9 overexpression activated the c-Myc-335 reporter (Fig. 2E). In combination with the ChIP data, these results were consistent with a model of SOX9 directly binding and activating this enhancer in conjunction with TCFs and β-catenin.

To understand how SOX9 was recruited to this enhancer, we searched its sequence for potential SOX9 binding sites. Our previous work on the *cis*-regulatory logic of c-Myc-335 had identified 4 TCF binding sites, 2 sites bound by CDX proteins, and 5 repeated motifs we named CAG sites, all of which contributed to enhancer activity (*9*). Using SOX9 binding site data from the JASPAR database, we scanned the c-Myc-335 sequence for putative clusters of SOX9 binding sites. This search identified a pair of putative SOX binding sites located adjacently in an inverted repeat orientation (Fig. 2F, S2A). The ability of SOX9 to dimerise and bind DNA is an important part of its ability to activate some of its transcriptional targets (*49*), and the binding sites in c-Myc-335 were in the correct orientation to be bound by a SOX9 dimer. We confirmed the ability of SOX9 to bind to these predicted sites using a gel-shift assay. A purified recombinant SOX9 fragment containing the DNA binding domain (SOX9 ^1-190^) induced a robust gel shift in a probe containing the WT SOX binding sites, and mutating either or both SOX9 binding sites in the probe attenuated the gel shift (Fig. 2G, S2B).

To functionally verify these SOX9 binding sites, we created a synthetic reporter containing multimerised copies of the SOX binding sites upstream of a minimal promoter. The activity of this reporter, named “SOXflash”, was SOX9-dependent, with reduced activity in LS174T cells upon RNAi-mediated SOX9 depletion (Fig. 2H). We then tested the contribution of SOX sites for WRE activation by mutating them in the c-Myc-335-luc reporter. Previously, we found that mutating the 4 TCF sites caused a reduction in enhancer activity (*9*). Mutating either one of the SOX sites caused a reduction in enhancer activity comparable to mutating the 4 TCF binding sites. Combined mutation of the TCF and SOX sites caused a nearly 200-fold reduction in enhancer activity (Fig. 2I).

### SOX9 is important for activating a subset of WREs in cancer cells

Our analysis of the c-Myc-335 WRE revealed that it was occupied by both TCFs and SOX9, and that its activity was dependent on both TCF and SOX9 binding sites. While the c-Myc-335 WRE is well-known for its role in CRC risk, the *MYC* gene is surrounded by several WREs implicated in the misregulation of *MYC* in many different kinds of cancer (*31*). Our examination of HT-29 SOX9 ChIP-seq data (*48*) identified multiple putative SOX9-bound enhancers around the *MYC* locus (Fig. 3A). Cross-referencing these regions with previously characterised WREs of *MYC*, we identified three WREs that overlap with SOX9-bound regions (Fig. 3B, S3A) (*31, 50*–*52*). Analogous to c-Myc-335, we named these regions Myc+8, Myc-29, and Myc-521, based on the distance in kilobases between the enhancers and the TSS of the *MYC* gene.

**Figure 3.**
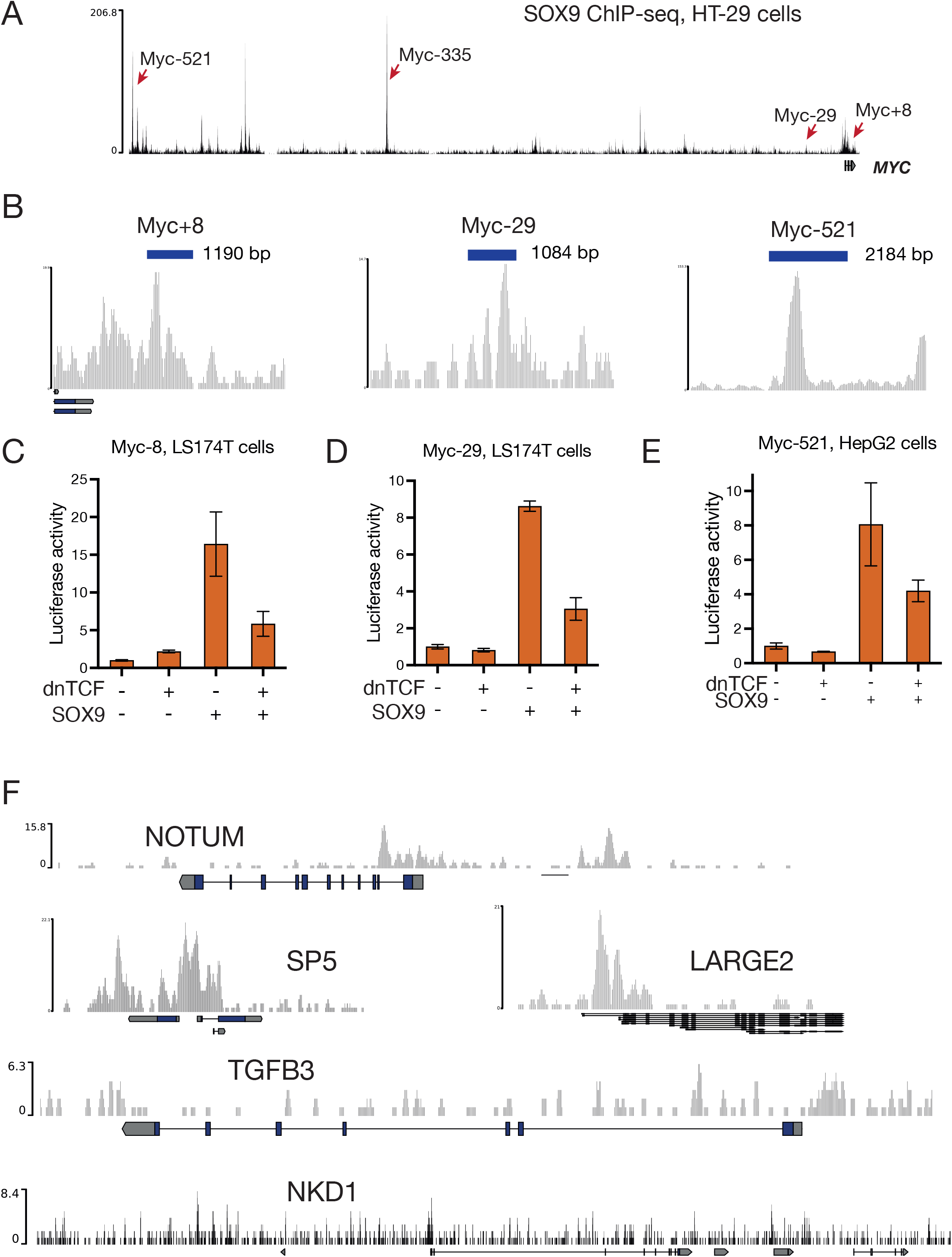
SOX9 and Wnt together regulates a multitude of WREs in cancer cells. **A)** Cartoon of the *MYC* gene locus showing the positions of cancer risk-associated WREs bound by SOX9 in HT29 cells. **B)** ChIP-seq tracks showing the binding of SOX9 to the Myc+8, Myc-29, and Myc-521 WREs. Dataset generated by (*48*). **C-E)** Luciferase assays showing the effect of overexpressing dnTCF, SOX9, or both on the activity of the Myc+8, Myc-29, and Myc-521 reporters. Indicated cell lines were transfected with reporters and protein expression constructs 24h before being assayed for luciferase activity. **F)** SOX9 ChIP-seq tracks of top 5 Wnt target genes in HT29 cells. HT29 RNA-seq data from (*54*) was cross-referenced with ChIP-seq data from (*48*).

To confirm that these enhancers were jointly regulated by Wnt signalling and SOX9, we generated luciferase reporters by cloning these regions upstream of a minimal promoter and a luciferase coding sequence. In LS174T cells, both Myc+8 and Myc-29 were activated by SOX9 overexpression, which was silenced by the overexpression of the Wnt pathway antagonist dnTCF (Fig. 3C,D). We then tested Myc-521 in HepG2 hepatocellular carcinoma cells which express a truncated form of β-catenin resulting in elevated Wnt signalling (*53*). Like Myc+8 and Myc-29, Myc-521 enhancer activity was stimulated by SOX9 overexpression and inhibited by additional dnTCF overexpression (Fig. 3E). These results suggest that SOX9 is a co-factor important for the activation of multiple *MYC* WREs in cancer cells.

Our findings raised the question of whether the positive role of SOX9 in WRE regulation was confined to *MYC*, or whether other Wnt target genes in CRC were also activated by SOX9. We cross-referenced the HT-29 SOX9 ChIP-seq dataset (*48*) with an RNA-seq dataset of Wnt-regulated genes in HT-29 cells (*54*). Of the top 5 high-confidence Wnt targets identified by this study, 3 genes (*NOTUM, SP5*, and *LARGE2*) had significant SOX9 peaks near their promoters and gene bodies, while no significant enrichment was seen near the other 2 (*NKD1, TGFB2*) (Fig. 3F). These findings suggest that SOX9 binds and regulates a subset of WREs in CRC cells.

### Synergistic activation of Paneth cell markers by Wnt signalling and SOX9

To identify WREs regulated by Wnt signalling and SOX9 in the context of normal development, we examined genes expressed specifically in Paneth cells, which require both Wnt signalling and SOX9 for their differentiation (*22, 25, 26*). Transcript levels of two human Paneth cell marker genes, *Defensin alpha 5* (*Defa5*) and *Defensin alpha 6* (*Defa6*) have been recently shown to be upregulated in CRC cells by SOX9 overexpression (*27*). The promoters of two human Paneth cell markers, *Defensin alpha 5* (*Defa5*) and *Defensin alpha 6* (*Defa6*) are known to contain multiple TCF binding sites, and transcriptional reporters made from these promoters are activated by Wnt signalling in cell culture (*28, 29*). We were interested in whether these Paneth cell WREs were also directly regulated by TCFs and SOX9.

To test whether *Defa5* was co-regulated by SOX9 and the Wnt pathway, we examined its expression in a HEK293 cell line with a DOX-inducible SOX9 overexpression construct (DOX-SOX9 cells) (*20*). We found that *Defa5* mRNA levels were synergistically upregulated under high-Wnt, high-SOX9 levels induced by DOX treatment in conjunction with the β-catenin destruction complex inhibitor CHIR-99021 (Fig. 4A). We then cloned the promoters of both *Defa5* and *Defa6* into the pGL4.10 promoterless luciferase reporter plasmid to generate transcriptional reporters for both genes (referred to as Defa5-luc and Defa6-luc). We tested the response of both reporters to SOX9 overexpression and Wnt pathway stimulation by the overexpression of a stabilised β-catenin (β-catenin*) (*55*). Both Defa5-luc and Defa6-luc were mildly activated by Wnt stimulation or SOX9 overexpression, but showed maximal activity under high-Wnt, high-SOX9 conditions similar to those found in Paneth cells (Fig. 4B,C). In LS174T cells which already have high levels of Wnt signalling, SOX9 overexpression activated Defa5 and Defa6 promoter activity, which was repressed by inhibiting Wnt signalling with dnTCF (Fig. 4D,E).

**Figure 4.**
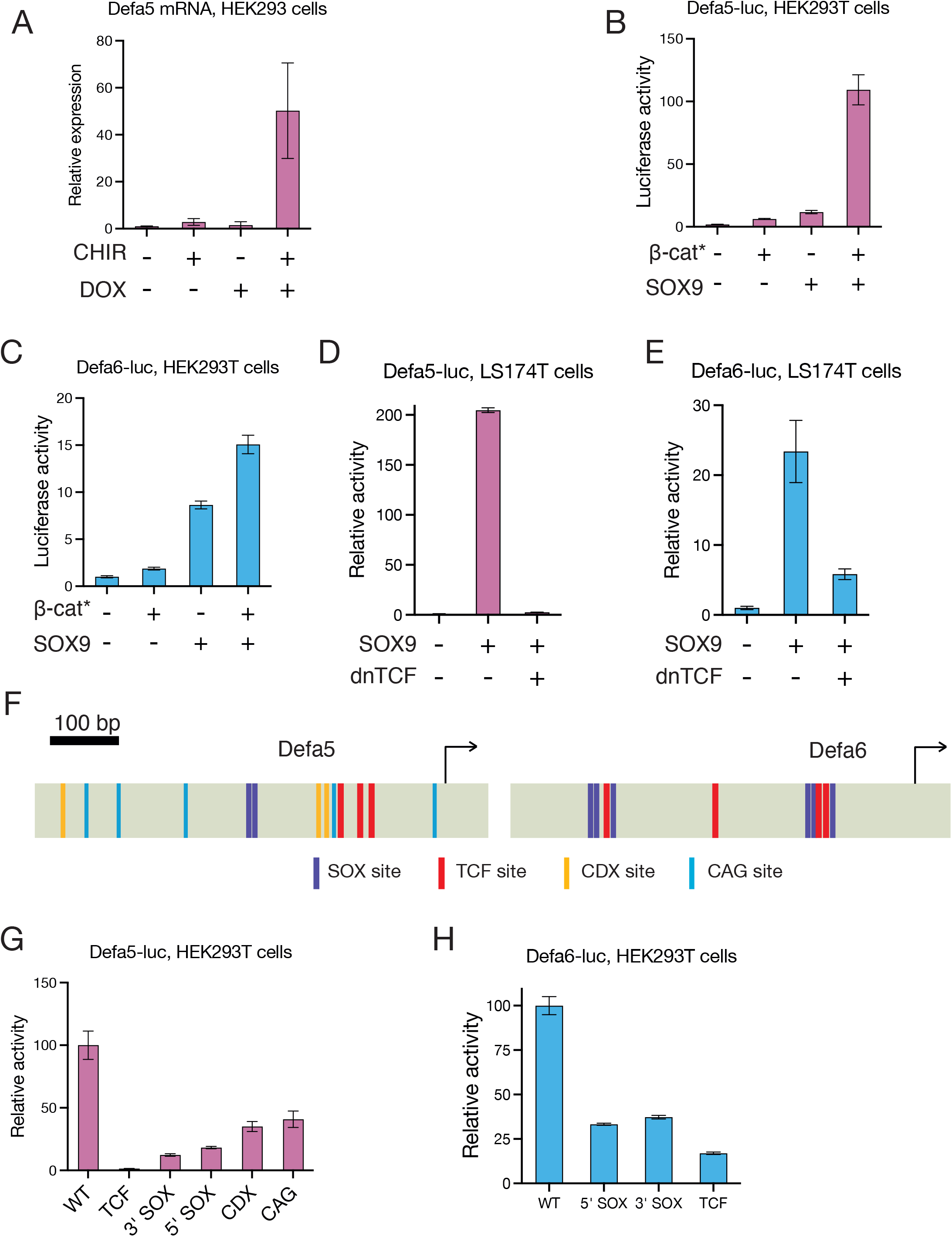
SOX9 and Wnt signalling synergistically drive Paneth cell defensin activity through TCF and SOX binding sites. **A)** RT-qPCR data showing induction of *Defa5* mRNA levels under high Wnt, high SOX9 levels. DOX-SOX9 HEK293 cells (*20*) were treated with CHIR-99021 to activate Wnt signalling and DOX to activate SOX9 overexpression. Transcript levels were measured 24h after treatment. **B)** Induction of Defa5-luc reporter activity by Wnt signalling and SOX9 in HEK293T cells. **C)** Induction of Defa6-luc reporter activity by Wnt signalling and SOX9 in HEK293T cells. **D)** Activation of Defa5-luc reporter activity by SOX9 overexpression and repression by dnTCF overexpression in LS174T cells. **E)** Activation of Defa6-luc reporter activity by SOX9 overexpression and repression by dnTCF overexpression in LS174T cells. **F)** Cartoons of the promoters of *Defa5* and *Defa6* showing the locations of TCF, SOX, CDX, and CAG sites. **G)** Loss of Defa5-luc reporter activity upon mutation of TCF, SOX, CDX, or CAG sites. **H)** Loss of Defa6-luc reporter activity upon mutation of TCF or SOX sites. For B-E, G, and H, cells were transfected with indicated reporters and plasmids expressing stabilised β-catenin (β-catenin*), Flag-SOX9, dnTCF, or empty vector (pcDNA3.1). Luciferase activity was measured 24h post transfection.

We then probed the *cis-*regulatory logic of the *Defa5/6* promoters. Computational searches for TF binding sites identified putative TCF and SOX binding sites in the promoters of *Defa5/6*, in addition to the CDX and CAG sites identified in several other WREs (Fig. 4F, S4A,B). We tested the importance of these sites by mutating the TCF, SOX, CDX, and CAG sites in *Defa5* and the TCF and SOX sites in *Defa6*. Measurements of reporter activity in HEK293T cells under high Wnt, high SOX9 conditions (Fig. 4G,H) and in LS174T cells with SOX9 overexpression (Fig. S4C, D) showed that all 4 sets of motifs were essential for WRE activity. Put together, these results indicate that in addition to working together to direct Paneth cell fate, Wnt signalling and SOX9 continue to be important after differentiation by directly activating genes expressed in Paneth cells using a regulatory logic similar to that seen in SOX9-activated oncogenic enhancers of *MYC*.

### TCF and SOX binding sites are sufficient for synergistic enhancer activation by Wnt signalling and SOX9

While we found no evidence of SOX9 acting as a Wnt antagonist in LS174T and DLD-1 cells, SOX9 is known to potently antagonise WRE expression in HEK293T cells (*20*), making the identification of WREs that were activated synergistically by Wnt signalling and SOX9 in HEK293T cells very striking. Previous work from our group characterised CREAX, a distal Wnt-responsive enhancer of the *Axin2* Wnt target gene and found that enhancer activity was mediated by a combination of TCF, CDX, and CAG sites (*9*). However, CREAX lacks SOX9 binding sites and is repressed by SOX9 overexpression (*20*). While the Defa5/6 and c-Myc-335 WREs also contain TCF, CDX, and CAG sites, they were set apart from CREAX by the presence of SOX9 binding sites. This suggested that the *cis-*regulatory grammar of these WREs was the sole determinant of whether they were activated or repressed by SOX9.

To directly test this idea, we generated a synthetic enhancer with a combination of TCF or SOX9 binding sites. Reporters containing only multimerised TCF binding sites (TOPflash) are robustly activated by Wnt signalling (*56*), and similar reporters containing SOX9 binding sites (SOXflash) show SOX9-dependent enhancer activity (Fig. 2H). Our new reporter, named TOP/SOX, contains 2 TCF and 2 pairs of SOX9 binding sites in the dimer-binding orientation (Fig. 5A, S5). In LS174T cells, TOP/SOX-luc was repressed by dnTCF and activated by SOX9 overexpression, mimicking the expression pattern of the c-Myc-335 enhancer (compare Fig. 5B and 2E).

**Figure 5.**
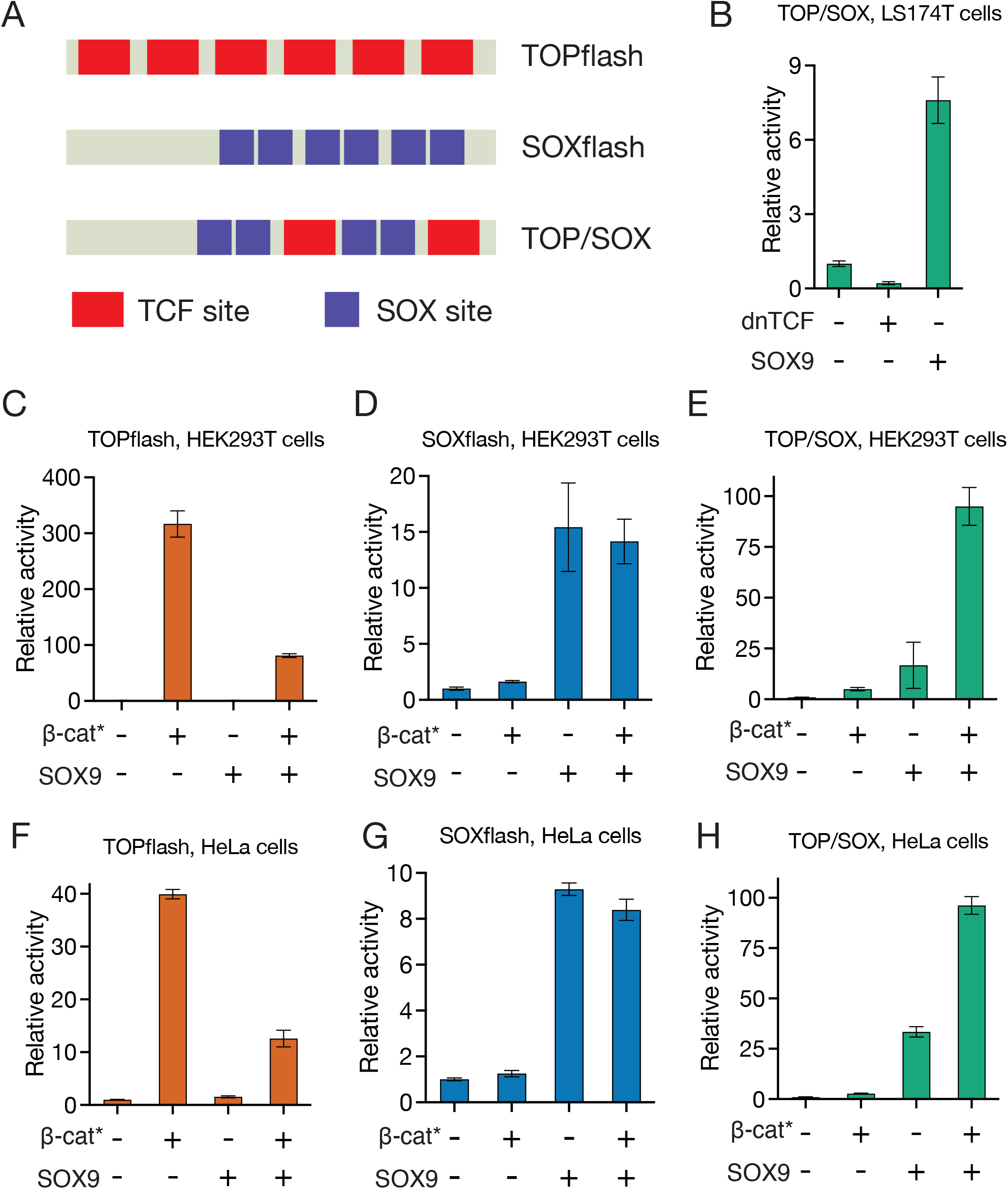
TCF and SOX binding sites are sufficient for synergistic enhancer activation by Wnt signalling and SOX9. **A)** Cartoons of synthetic enhancers containing combinations of TCF and SOX binding sites. TOPflash contains only TCF sites, SOXflash contains only SOX sites, and TOP/SOX contains both TCF and SOX sites. **B)** TOP/SOX is repressed by dnTCF overexpression and activated by SOX9 overexpression in LS174T cells. **C-H)** Relative activity levels of TOPflash, SOXflash, and TOP/SOX in HEK293T (C-E), and HeLa cells (F-H) upon stimulation of Wnt signalling, SOX9 overexpression, or both. Cells were transfected with indicated reporters and overexpression constructs, and luciferase activity was assayed 24h later.

We then compared the response of TOPflash, SOXflash, and TOP/SOX in HEK293T and HeLa cells to the overexpression of β-catenin*, SOX9, or both. As expected, TOPflash activity was robustly induced by β-catenin* and repressed by SOX9 overepxresion (Fig. 5C). In contrast, SOXflash was robustly activated by SOX9 overexpression and not significantly affected by β-catenin* (Fig. 5D). Strikingly, TOP/SOX was maximally activated by the combined overexpression of both β-catenin* and SOX9 (Fig. 5E). A similar pattern of expression was seen in HeLa cells, with TOP/SOX showing a qualitatively different expression pattern to TOPflash and SOXflash (Fig. 5F-H). The conditions that cause dramatic reductions in TOPflash activity are identical to those that result in maximal activation of TOP/SOX (Fig. 5C,F). Our data demonstrate that within a single cell type, the presence or absence of SOX9 binding sites can determine whether a WRE is repressed or activated by SOX9.

### TCFs and SOX9 directly interact through non-DNA contacting residues in their DNA binding domains

Multiple SOX family members have been shown to complex with TCFs through their DNA-binding HMG domains (*10*), and we wondered if the same was true in the case of SOX9. We tested the ability of purified recombinant GST-tagged LEF1 and TCF7 to interact with His-tagged SOX9 in a pulldown assay, with GST as a negative control. Full-length SOX9 directly bound to both LEF1 and TCF7 *in vitro* (Fig. 6A,B). A GST-tagged fragment of LEF1 containing just the HMG domain (Fig. 6A) also robustly pulled down full-length SOX9 (Fig. 6C), suggesting that like other TCF-SOX protein interactions, TCF-SOX9 interactions were also mediated by HMG domains. To eliminate the possibility that this interaction was scaffolded by residual bacterial DNA in our purified protein preparations, we tested the impact of Mnase treatment, and found no effect on the interaction between the HMG domains of LEF1 and SOX9 (Fig. S6A). This confirmed that the HMG domain of SOX9 could directly interact with the HMG domain of TCFs, independent of DNA binding.

**Figure 6.**
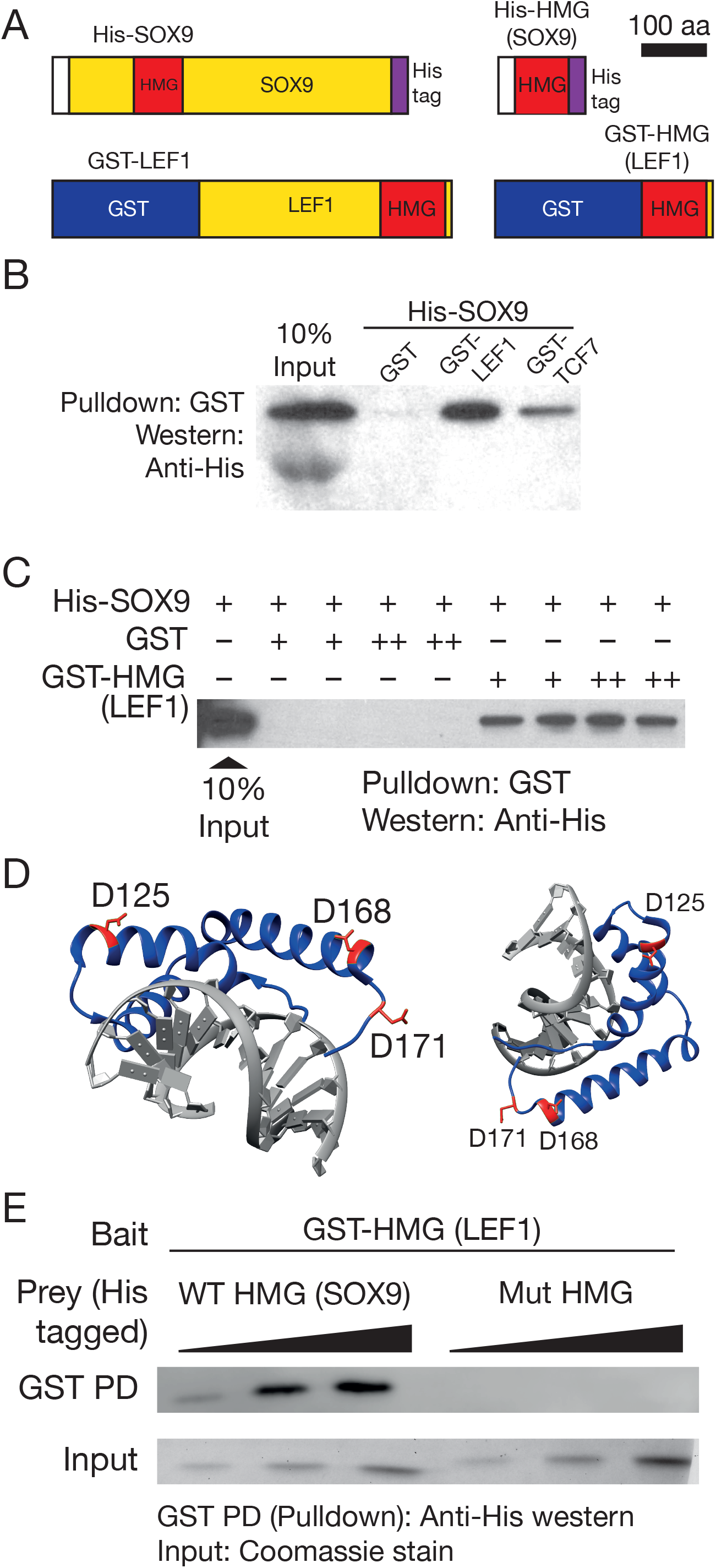
SOX9 directly binds to TCFs through non-DNA contacting residues in its DNA-binding HMG domain. **A)** Cartoons of GST-tagged TCF7, LEF1, and LEF1 fragments used for pulldown experiments. **B)** GST pulldown showing direct binding of SOX9 to LEF1 and TCF7. **C)** GST pulldown showing binding of full-length SOX9 to the HMG domain of LEF1. **D)** Two views of a crystal structure of the SOX9 HMG domain (PDB ID: 4EUW). Amino acid residues mutated to abolish TCF binding are coloured red with their side chains shown. **E)** Mutation of 3 amino acids in SOX9’s HMG domain (D125, D168, and D171) abolishes the TCF-SOX interaction. Purified His-tagged proteins were pulled down using glutathione beads with indicated GST-tagged proteins as bait for B, C, and E. GST was used as negative control.

To test the importance of TCF/SOX9 interactions in gene regulation, we wanted to generate a separation-of-function mutant of SOX9 that retained its ability to bind to DNA but could not bind to SOX9. An analysis of a crystal structure of the HMG domain of SOX9 complexed with DNA (PDB ID: 4EUW) revealed the presence of multiple charged residues on the outer surface of the HMG domain. Working off the hypothesis that the TCF/SOX9 interaction was mediated by charged residues whose side-chains pointed away from the DNA, we tested constructs containing charge swap mutations for their ability to bind to the HMG domain of LEF1. We found that a SOX9 HMG domain fragment containing mutations changing three aspartate (acidic) residues in SOX9 (D125, D168, and D171) to lysines (basic) showed no detectable binding to the HMG domain of LEF1 (Fig. 6D,E). These results suggested that TCFs and SOX9 bound to each other through salt bridges mediated by non-DNA contacting residues on their HMG domains. They also provided us with a reagent that could be used to test the role of TCF/SOX9 protein-protein interactions in gene regulation. The interacting residues were highly conserved across vertebrate orthologs of SOX9 (Fig. S6B), suggesting that TCF-SOX9 interactions may be important for gene regulation across many species. Within human SOX proteins, the three residues were highly conserved (identical or conserved mutation) in the SOXE (SOX8,9,10), and SOXC (SOX4,11,12) subfamilies, with limited conservation in other family members (Fig. S7).

### TCF/SOX9 interactions are essential for synergistic enhancer activation and cancer cell survival

Before using our novel mutant to test the importance of TCF/SOX9 interactions for gene regulation, we wanted to ensure that the mutations did not compromise the ability of SOX9 to perform its TCF-independent functions. First, we compared the ability of WT and D125,168,171K Mutant (Mut) SOX9 to activate the SOXflash reporter in HEK293T cells. Transfection of identical amounts of plasmid expressing WT or Mut SOX9 produced similar activation of SOXflash (Fig. 7A). The ability of SOX9 to repress Wnt reporters such as TOPflash involves the transcriptional activation of the SOX9 target gene MAML2 (*20*). In HEK293T cells, both WT and Mutant suppressed β-catenin*-induced TOPflash activation to similar levels (Fig 7B). These experiments suggested that the ability of SOX9 to enter the nucleus, complex with DNA, and mediate transcriptional activation was not affected by the HMG domain mutations.

**Figure 7.**
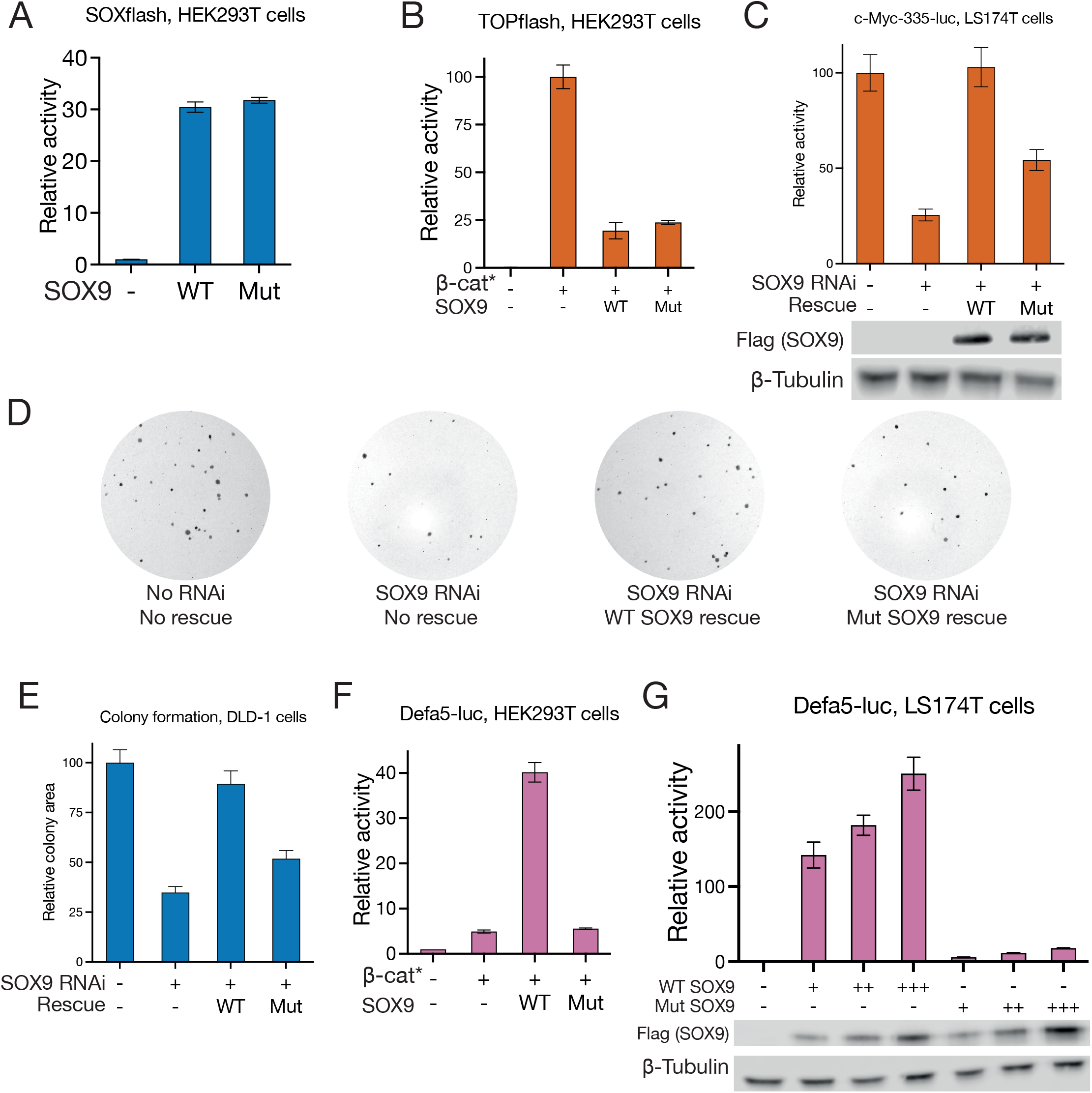
TCF/SOX interactions are essential for WRE activity and colorectal cancer cell growth. **A)** Luciferase assay showing that SOXflash activity can be driven by both WT and TCF interaction-deficient (Mut) SOX9. **B)** Luciferase assay showing both WT and Mut SOX9 are equally capable of suppressing TOPflash activation by β-catenin*. **C)** Luciferase assay showing that WT SOX9, but not Mut can rescue the loss of c-Myc-335 reporter activity caused by SOX9 RNAi in LS174T cells. Western blots show relative expression levels of WT and Mut SOX9. **D)** Representative images showing that WT SOX9, but not Mut can rescue the loss of colony formation caused by SOX9 RNAi in DLD-1 cells. **E)** Quantification of the area occupied by colonies in (D). **F-H)** Luciferase assays showing that Defa5-luc and Defa6-luc can be activated by WT but not Mut SOX9 in HEK293T and LS174T cells. Western blots in (H) show relative expression levels of WT and Mut SOX9.

We then compared the ability of WT and Mut SOX9 to drive c-Myc-335 enhancer activity. While the overexpression of WT SOX9 could rescue the loss of reporter activity caused by SOX9 RNAi, overexpression of Mut SOX9 at similar levels only resulted in a partial rescue (Fig. 7C). Similarly, overexpression of WT but not Mut SOX9 rescued the colony formation defect in DLD-1 cells caused by SOX9 RNAi (Fig. 7D,E). In the case of Defa5-luc, the overexpression of Mut SOX9 along with β-catenin* failed to produce the synergistic activation shown by WT SOX9 in HEK293T cells (Fig. 7F). In LS174T cells, overexpression of Mut SOX9 produced a very attenuated activation of the Defa5-luc reporter when compared to similar levels of WT SOX9 overexpression (Fig. 7G). Finally, Mut SOX9 also showed a compromised ability to activate the Defa6-luc reporter in concert with β-catenin* in HEK293T cells (Fig. S8A), and its overexpression in LS174T cells resulted in reduced Defa6-luc reporter activation when compared to WT SOX9 (Fig. S8B). Put together, these results suggest that a combination of cis-regulatory grammar and TCF/SOX9 protein-protein interactions is essential for activating a Wnt/SOX9 target gene program that allows Paneth cell gene expression and the growth and survival of CRC cells.

## Discussion

How the interplay between signalling-dependent developmental regulators like the Wnt pathway and lineage-determining transcription factors such as SOX proteins results in spatio-temporally diverse patterns of gene expression is a fundamental question in developmental biology. Prior to this study, models of SOX9’s role in Wnt signalling characterised SOX9 as a Wnt antagonist, primarily influenced by work on the mammalian sex determination pathway and cell culture studies that examined the effects of SOX9 overexpression on Wnt signalling (*10*). In conjunction with several recent studies (*27, 37, 38*), our findings demonstrate a highly context-specific role for SOX9 in CRC, in which it promotes the growth and survival of CRC cells (Fig. 1). This transcriptional program involves SOX9 directly binding and regulating a subset of WREs (Figs. 2, 3). Findings of SOX9 activating Paneth cell markers suggest a role for Wnt signalling and SOX9 cooperating to activate gene expression during both normal and cancerous growth (Fig. 4). Whether SOX9 activates or represses a given WRE is determined by the presence or absence of SOX9-binding sites (Fig. 5), and the ability of SOX9 to complex with TCFs is integral to its ability to activate Wnt targets (Figs. 6, 7).

Previous work on SOX9 identified a naturally occurring shorter isoform of SOX9, named miniSOX9, that was expressed in many CRC lines (*57*). When overexpressed, miniSOX9 inhibited SOX9-driven reporter activity and also alleviated SOX9’s inhibition of Wnt readouts, leading to a model of miniSOX9 as a naturally occurring dominant negative variant that allowed the *SOX9* gene to stimulate cancer growth by inhibiting the ability of full-length SOX9 to repress Wnt signalling (*57*). However, this model was not tested directly with loss-of-function experiments. Our data show that SOX9 is not be a potent Wnt antagonist in CRC cells at endogenous levels of expression (Fig. 1A, S1A). Additionally, overexpression of full-length SOX9 was sufficient to rescue the loss of colony formation after SOX9 KD (Fig. 7D,E) targeting the 5’-UTR that is shared by full-length and WT SOX9 (*57*). These results establish a role for the full-length SOX9 protein in promoting cancer growth by transcriptional activation.

Our examination of the regulatory logic of SOX9-activated WREs revealed the presence of functionally important TCF and SOX9 binding sites without any easily discernible binding site grammar (Fig. 2,4). While the validated TCF and SOX binding sites in c-Myc-335 and the *Defa5* promoter are separated by several dozen nucleotides, they are more tightly clustered in the *Defa6* promoter (Fig. 2F,4F). Similarly, the predicted TCF and SOX9 binding sites in Myc+8, Myc-29, and Myc-521 also do not follow a readily observable pattern of binding site grammar. Together with our synthetic reporter experiments (Fig. 5), this suggests that the mere presence or absence of SOX binding sites is sufficient to discriminate between activation and repression of WREs by SOX9. While some of the SOX9 binding sites characterised in this study are in the correct orientation to be bound by SOX9 dimers, others are singular binding sites. Previous work has shown that the ability of SOX9 to dimerise is crucial for its role in some contexts such as chondrogenesis, but not in others such as the mammalian sex determination pathway (*49, 58*). In this study, we only tested the effect of individual SOX binding site mutations in the context of c-Myc-335 and *Defa5*, which have a single pair of SOX9 binding sites each. While this data might indicate a role for SOX9 dimerisation in Wnt target gene regulation, more mutagenesis experiments and experiments using dimerization-deficient mutants are required to test it directly.

Previous work has shown that when overexpressed, SOX9 inhibits WREs in a dose-dependent manner. SOX9 overexpression results in the transcriptional upregulation of *MAML2*, which promotes β-catenin degradation (*20*). At the same time, overexpression of SOX9 in CRC cells with high levels of Wnt signalling activates a Paneth cell-like gene expression program (*27*). While intestinal stem cells retain their ability to proliferate and differentiate in the absence of SOX9 (*25, 26*), they do show changes in their gene expression patterns after SOX9 removal (*59*). The requirement of SOX9 for proliferation in these cell types has led to the idea that a “critical dose” of SOX9 may be important for intestinal stem and cancer cell proliferation (*36, 60*). Our findings support this model and provide a molecular basis for it. The ability of SOX9 to promote the expression of a subset of Wnt target genes makes it an essential regulator of growth and proliferation in these cells. Genes co-regulated by Wnt signalling and SOX9 may also show differential sensitivity to SOX9 levels. For example, the c-Myc-335 enhancer showed high activity at the endogenous SOX9 levels seen in LS174T CRC cells and was only modestly activated by SOX9 overexpression (Fig. 2E). In contrast, the promoters of *Defa5/6*, which are active in Paneth cells that express higher levels of SOX9, were significantly activated by SOX9 overexpression (Fig. 4D). Our finding that Wnt/SOX9-activated enhancers are most active under conditions that lead to a partial inhibition of WREs lacking SOX9 binding sites (Fig. 5) may provide a mechanistic explanation for the “critical dose” model, with higher levels of SOX9 shutting down WRE activity through β-catenin degradation, and lower levels of SOX9 preventing the activation of some WREs, leading to reduced proliferation under both reduced and increased SOX9 levels.

While hyperactivated Wnt signalling has long been known as a driver of CRC, its important role in driving the homeostasis of many different adult tissues has proved an important roadblock for the use of Wnt pathway inhibition as a cancer therapy (*1, 30, 61*– *63*). Our identification of the importance of SOX9-TCF interactions as a driving force for Wnt target gene activation in CRC may allow the development of novel therapeutics for CRC that disrupt the TCF-SOX9 interaction. Short peptides that mimic TF-TF interaction domains have been used in the past to successfully and specifically disrupt TF binding *in vivo* (*64*). Similar peptides may allow the specific inhibition of Wnt signalling in cancer cells without depleting the many Wnt-dependent stem cell populations found in the body.

## Materials and methods

### Cell culture and transfection

Cell lines were grown at a temperature of 37⍰ with 5% CO_2_. HEK293T (ATCC, CRL-3216) and DLD-1 (ATCC, CCL-221) cells were grown in DMEM (Dulbecco’s Modified Eagle Medium, Gibco, 11995065) supplemented with 10% foetal bovine serum (FBS) and penicillin/streptomycin/glutamine (PSG, Gibco, 10378016). HepG2 cells were a kind gift from Dr. Jun Wu (University of Michigan). The LS174T-derived pTer-β-catenin-RNAi stable line (pTer-β-cat) was a kind gift from Dr. Xi He (Boston Children’s Hospital, Harvard Medical School. LS174T cells (ATCC, CL-188), their derivatives, and HepG2 cells were grown in MEM (Minimum Essential Medium, Gibco, 11095080) with 10% FBS and PSG. Tetracycline-screened FBS (Cytiva, SH30071.03T) was used to culture pTer-β-cat cells and their derivatives. pTer-β-cat cell media was supplemented with 100 µg/ml Zeocin (InvivoGen, ant-zn-05) to select for the β-catenin RNAi cassette.

Lentiviral supernatants were produced at the University of Michigan Vector Core. To generate cells stably expressing non-targeting (scrambled) or SOX9-targeting shRNAs (details in Table S1), cells were incubated with viral supernatants overnight. Cells were thereafter maintained in 1 µg/ml Puromycin to select for the Scrambled or SOX9 shRNA expressing cassette.

Transient transfections of HEK293T, HepG2, and LS174T cells done with PEI MAX (Polysciences, 24765-1). DLD-1 cells were transfected with Lipofectamine 2000 (Invitrogen, 11668030) according to manufacturer’s instructions.

## Plasmids

SOX9 RNAi plasmids used to generate stable LS174T cell lines were from the pGIPZ library and purchased from the University of Michigan Vector Core. shRNA constructs used for transient transfection experiments were cloned into the pSUPER vector (Oligoengine, VEC-pBS-0002). Empty pSUPER plasmid was used as negative control. The c-myc-335 locus was amplified via PCR using human Jurkat cell genomic DNA as a template and cloned into XhoI/BglII sites in pGL4.23 (Promega, E8411). dnTCF (*65*) and Flag-SOX9 (*20*) constructs have been described previously. TF binding site mutations in c-Myc-335 were generated using a combination of Gibson assembly and site-directed PCR mutagenesis. Details of mutations are specified in Table S2. Myc+8, Myc-29, and Myc-521 reporters were amplified from HEK293T cell genomic DNA and cloned into pGL4.23 between the XhoI and HindIII restriction sites. The sequence of the region between these sites is shown in Fig. S3. Human β-catenin* containing the S33Y mutation was cloned into the pcDNA3.1 vector and was a gift from Dr. Eric Fearon, University of Michigan. *Defa5/6* reporters were PCR amplified from human genomic DNA and cloned into the pGL4.10 promoterless luciferase vector (Promega, E6651) between the XhoI and KpnI resctriction sites. gBlock gragments containing mutations in TF binding sites were ordered from IDT and cloned into these plasmids to generate site mutant reporters. Oligonucleotides containing fragments of the TOP/SOX reporter sequence were ordered from IDT, annealed together, and cloned into the pGL4.23 vector. The sequence of the synthetic reporter is shown in Fig. S5. GST-tagged TCF7 has been previously described (*9*). GST-tagged LEF1 was generated similarly by cloning the LEF1 ORF between the BamHI/NotI sites of pGEX-6p-1. His-tagged SOX9 was generated by cloning the ORF into the BamHI/SalI sites of pET-52b. HMG domain fragments of LEF1 and SOX9 were PCR amplified and cloned into pGEX-6p-1 and pET-52b between the same restriction sites used to make the full-length expression constructs. D125,168,171K mutations were introduced in the His-HMG and Flag-SOX9 expression plasmids using PCR with mismatched primers followed by Gibson assembly.

### Colony formation assays

Cells were plated in 12-well plates and transfected or treated with doxycycline for the indicated amounts of time. Following this, they were resuspended in the appropriate cell culture media after treatment with 0.25% Trypsin-EDTA. Cell numbers were determined for each condition using a haemocytometer. Cells were then diluted to obtain the final concentrations reported in each experiment and then seeded in triplicates onto 6-well plates. They were allowed to grow undisturbed for 12-14 days until colonies were easily visible to the naked eye. Colonies were fixed with methanol for 20 minutes, following which they were stained with crystal violet staining solution (0.5% (w/v) crystal violet, 20% methanol) for 10 minutes. They were then rinsed with distilled water, dried overnight, and imaged.

The area occupied by the colonies was quantified using Fiji (*66*). Each well was defined as a circular region of interest and then cropped out. Wells were individually thresholded and the fraction of the area occupied by the wells was measured.

### MTT assay for cell number

MTT assays were carried out using Cell Proliferation Kit I (Roche, 11465007001). Cells expressing scrambled or SOX9 shRNAs were plated onto 48-well plates with and without DOX at a density of 30,000 cells/well. Serial dilutions of cells were plated to determine a standard curve. 48 hours later, 20 µl of MTT labelling reagent was added to each well. 200 µl of solubilisation solution was added 4 hours later. Absorbance was measured at 565 nm using a Tecan Infinite 200 plate reader. Relative cell densities of the different samples and treatments were then calculated based on the standard curve. All measurements were done with biological triplicates.

### RT-qPCR for transcript quantification

RNA was extracted using the Rneasy mini kit (Qiagen, 74104). Reverse transcription was then performed using Superscript III Reverse Transcriptase (Invitrogen, 18080093). The Power SYBR Green PCR master mix (Applied Biosystems, 4368577) was used to perform PCRs using a CFX Connect Real-Time PCR Detection System (Bio-Rad, 1855201). Mean and standard deviations of transcript levels were quantified using the delta-delta Ct method using *G6PD* transcript levels for normalisation. All measurements were done with biological triplicates. Primers are shown in Table S4.

### Chromatin immunoprecipitation (ChIP)

TCF7L2 ChIP was performed and signals were quantified using qPCR as previously described (*9*). SOX9 ChIP was done in a similar fashion using an anti-SOX9 antibody (Sigma, AB5535). For β-catenin ChIP, cells were first treated with the protein-protein crosslinker EGS (ethylene glycol bis(succinimidyl succinate)) (Thermo Scientific, 21565) at a concentration of 12.5 mM (*65*), following which ChIP was performed using an antibody targeting active β-catenin (CST, 8814S). One set of primers for qPCR (Table S4) were designed to amplify inside the c-Myc-335 enhancer region and 2 primer pairs were designed to amplify flanking regions outside the enhancer for comparison. ChIP signals from each of these primers was compared to the signal from a control region to quantify enrichment. Primers used for ChIP are shown in Table S5.

### ChIP-seq data analysis

ChIP-seq data analysed in this study was generated by a previous study (*48*). Publicly available data was downloaded from GEO (Accession number:GSE63629). The data file GSM1554224_HT29_SOX9_antibody.bar was converted into bedgraph format using USeq (*67*). ChIP-seq track visualisations were generated using pyGenomeTracks (*68*).

### Computational analysis of transcription factor binding sites

TCF, CDX, and CAG sites were computationally annotated using FIMO (*69*) as previously described (*9*). Initial identification of SOX9 binding sites was based on the SOX9 MA0077.1 motif file from the JASPAR database (*70*). To ensure the identification of potential SOX9 dimeric binding sites, searches were repeated with lowered thresholds to look for lower-scoring sites in the correct orientation located adjacent to high-scoring sites.

### Luciferase assays and analysis

Luciferase assays were done with firefly-luciferase reporter plasmids and constitutively expressed renilla luciferase as an internal transfection control. Luciferase activity was measured using Promega’s Dual Luciferase Assay system (Promega, E1910) according to manufacturer’s protocols. Firefly/renilla ratio was used as the measure of activity for each well. All assays were done in triplicates, following which the mean and s.d. of firefly/renilla ratios were calculated for each experimental condition. To calculate relative luciferase activity (RLA), a basal condition was selected (specified in each figure) and the ratios of all conditions were expressed in proportion to it.

### Electrophoretic Mobility Shift Assay (EMSA)

Binding reactions were carried out in Binding Buffer (10 mM Tris-Cl pH 7.5, 5 mM MgCl _2_, 50 mM KCl, 1 mM DTT, 2.5% glycerol, and 0.5% NP-40), 50 ng/mL poly dI·dC, 40 fmol biotinylated oligonucleotides (IDT). The reactions were then allowed to sit on ice for 5 minutes then at room temperature for 15 minutes. The reactions were then loaded onto a pre-run 6% TBE polyacrylamide gel followed by transfer to a Biodyne B positively charged membrane (Pall life sciences, 60208). Membranes were developed using a Light Shift chemiluminescent EMSA kit (Thermo Scientific, 20148).

### Recombinant protein expression and purification

GST and His-tagged protein expression constructs were expressed and purified as previously described (*9*). Protein expression was induced with 1 mM IPTG and cells were collected 3-4 h later. All constructs except for the His-tagged mutant SOX9 HMG were grown using the BL21(DE3) *E. coli* strain (Thermo Scientific, EC0114). The mutant HMG construct was expressed and purified in the Rosetta(DE3) strain (Novagen, 70954).

### GST pulldown assays

The concentration of purified recombinant proteins was measured using Bradford’s reagent (Bio-Rad, 5000202). 0.25-2 µg of each protein was used in a binding reaction. Binding reactions were carried out in a binding buffer containing 25 mM HEPES pH 7.5, 12.5 mM MgCl2, 300 mM KCl, 0.1% NP-40, 1 mg/mL BSA, 1 mM DTT. GST and His-tagged proteins were mixed in binding buffer to a total volume of 200 µL and incubated in a rotator for 1h at 4°C. Glutathione-sepahrose (Thermo-Scientific, 25236) were washed thrice in binding buffer and then added to binding reactions at a packed bead volume of 10 µl per reaction. After another 1h incubation at 4°C with rotation, beads were washed thrice in washing buffer (25 mM HEPES pH 7.5, 12.5 mM MgCl2, 400 mM KCl, 0.5% NP-40) and eluted by boiling for 5 minutes in 2x SDS buffer (100 mM Tris-Cl pH 6.8, 4% SDS, 0.2% bromophenol blue, 20% glycerol). Samples were then analysed by SDS-PAGE followed by western blotting.

### Western blotting

Western blotting was performed after SDS-PAGE as previously described (*9*). Western blots were probed with the following antibodies: Anti-Flag-HRP (Sigma-Aldrich, A8592), Anti-SOX9 (Sigma-Aldrich, AB5535), Anti-β-catenin (BD Biosciences, 610154), Anti-β-Tubulin (Proteintech, 66240-1-Ig), Anti-His (Cytiva, 27-4710-01).

### Protein structure visualisation

Protein structures were downloaded from RSC PDB and visualized with UCSF ChimeraX (*71*), developed by the Resource for Biocomputing, Visualization, and Informatics at the University of California, San Francisco, with support from National Institutes of Health R01-GM129325 and the Office of Cyber Infrastructure and Computational Biology, National Institute of Allergy and Infectious Diseases.

## Acknowledgements

The authors would like to thank Haidar A. Haidar and Dr. Abhishek Sinha for contributing to the initial stages of this project, and Aarti Kejriwal and Matthew Benson for generating some of the constructs used in this study.

## Conflict of interest

The authors declare that they have no conflicts of interest.

## Funding

This work was supported by grants from the US National Institute of Health [R01 GM108468 to K.M.C]; University of Michigan Rogel Cancer Center [to K.M.C]; University of Michigan M-cubed [to K.M.C]; University of Michigan Department of Molecular, Cellular, and Developmental Biology [Biosciences fellowship to A-B.R.]; and the Rackham Graduate School, University of Michigan [One-Term Dissertation Fellowship and a Graduate Student Research Grant to A-B.R.].

## Notes

### Competing Interest Statement

The authors have declared no competing interest.

