## Supplemental figures for "SOX9 binds TCFs to mediate Wnt/β-catenin target gene activation"

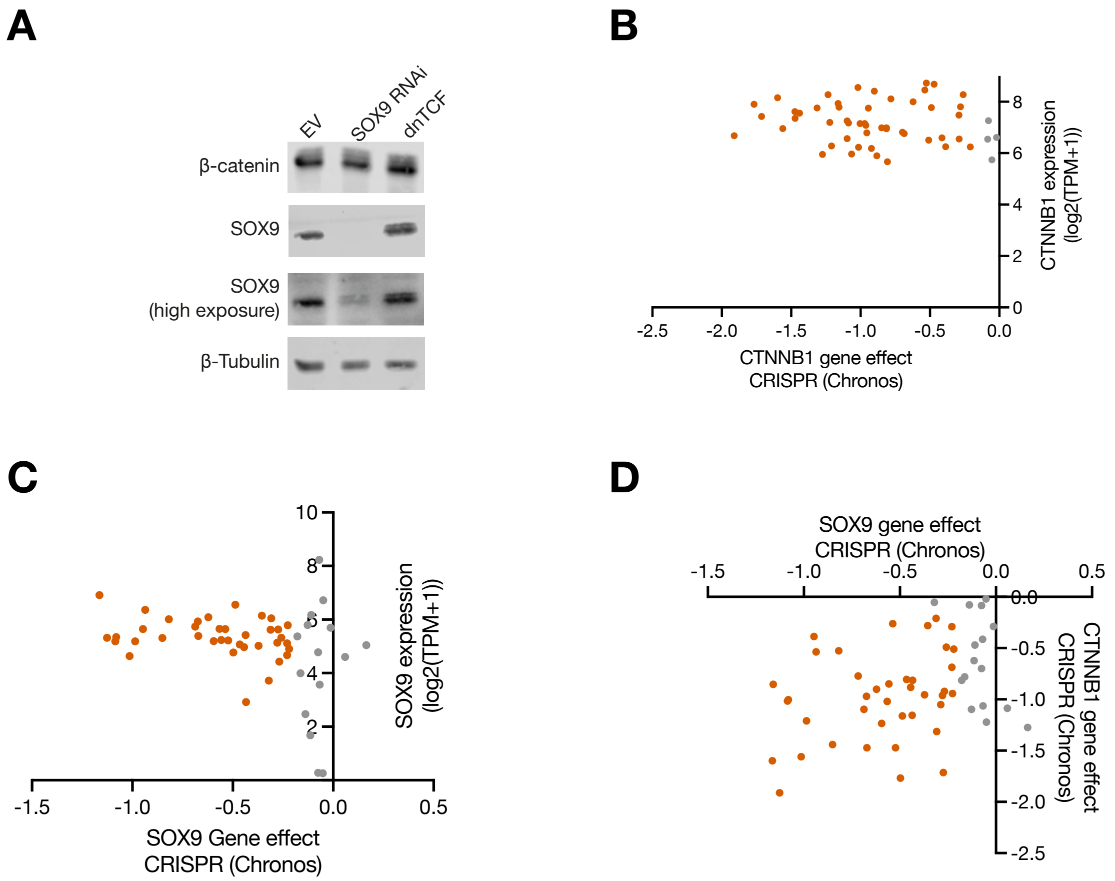


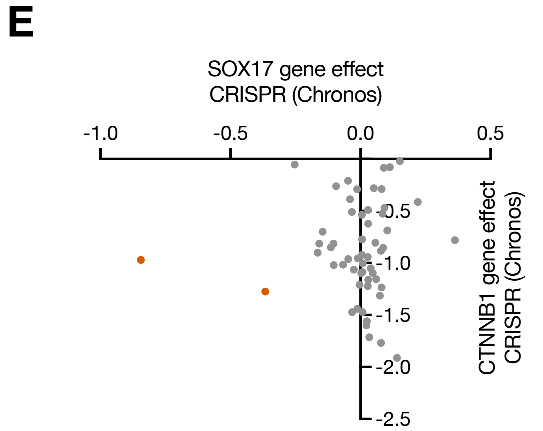


**F**

| Gene | Entrez Id | Correlation |
|---|---|---|
| **SOX10** | 6663 | 0.5211 |
| **DUSP4** | 1846 | 0.3923 |
| **BRAF** | 673 | 0.3753 |
| **MAPK1** | 5594 | 0.3561 |
| **CYB5R4** | 51167 | 0.3066 |
| **MITF** | 4286 | 0.2945 |
| **CTNNB1** | 1499 | 0.2928 |
| **MAP2K1** | 5604 | 0.2451 |
| **FASN** | 2194 | 0.2312 |
| **NEDD9** | 4739 | 0.2299 |

**Fig S1. SOX9 is a regulator of CRC cell growth and proliferation.**
**A)** Western blots showing no change in β-catenin levels upon SOX9 RNAi and no change in SOX9 levels upon inhibition of Wnt signalling by dnTCF overexpression in DLD-1 cells. Cells were transfected with either the empty pcDNA3.1 vector (EV), or plasmids expressing an shRNA targeting SOX9 or a dnTCF (dominant negative TCF) overexpression construct and harvested for western blotting 48h later. **B)** Correlation between CTNNB1 expression and essentiality in 53 Cancer Cell Line Encyclopedia (CCLE) CRC lines. **C)** Correlation between SOX9 expression and essentiality in 53 CCLE CRC lines. **D)** Correlation between essentiality scores of SOX9 and CTNNB1 (β-catenin) in 56 CCLE CRC lines. **E)** Correlation between SOX17 gene effect and CTNNB1 gene effect in 56 CCEL CRC lines. In **(B-E)**, essentiality (gene effect) is measured using Chronos scores. Each data point represents a single cell line. Cell lines passing threshold Chronos score values of -0.2 and expression values of 2 TPM are coloured orange, while others are coloured grey. All data were sourced from the 22Q1 public DepMap release. **F)** Top 10 genes with essentiality scores most correlated with SOX9 across all CCEL cancer cell line datasets. β-catenin shows the 7$^{th}$ best correlation.

## A

c-Myc-335 enhancer sequence:
Chr8:127400835–127401814

CGCTCCATAGAGCCTGCAGAGGGCACTAGACTGGGAATTAGAAAACCTGATTTCCCTTCCAGCTCCAC
CTCTGACCAATTGCCTGACCCTGGTCAAATTGCTTAACCTCTTCCTATCTCAGCTCCCTATCCATAAA
ACAGAGGGACGAATAAACTCTCCTCCTACCACTAAGAGGTGTAGCCAGAGTTAATACCCTCATCGT**CC**
**TTTGAGC**TCAGCA**GATGAAAGG**CACTGAGAAAA**GTACAAAGA**ATTTTTATGTGCTATTGACTTTATTT
TATTTTATGTGGGGGAGGGAGCCGGCCCCAGCTGGAAAGCTGCTTTCTCTG**AATCAAAGG**GCAGGAAC
CCAGCAAGTTTCTCAGGATTGGGGCCTTAGACTGGGCTGTGTATACAGACAGTGCCAGCCAACCCCAC
AGTTCAGTTTCCTTTAACCTGGTGCTCCAGGCAATAACTGTGCAACTCTGCAATTT**AACAATGT**GT**TC**
**TTTGTC**CCACAACTGTTCTCGTTTCTCAACTGCCCAGGTAATATGTTTGGGCCTGTAGGAAGAGTCAA
ATAGTTAATAAGGGAAGGGTTTGGCATGCCCTACGTAAGTTCTACCAGCAAGTCCCAACAAGAAGGCA
TTCTGTGTCTCCTGATTCCTGACCTACCCCCAAAATGTACAAATGTACAAGGAATGAGCCCACTTTCC
CAGCAGGCTGTAATACCAGTTTGGCCTATATCAATGCATTGGTGAGCTGTGTTTTGTTTATGGTTTTA
TGCCATCTATTTTCCCATGGATATTATGTTTTCTAAAGAGCCCTTAAGTTTACGTCAGCTTTTAAAGC
TACCAGCAGCACCATTTCAGTTCATATTAAGCCCTTAATATGGTATGAATAGGAGAGCTATTAGACTA
AAGAGCCATAATCATCCCTGAGGAAAACATCCATCACCAACATTTATGTGGTCCCTGAACTTCTAAAA
GGTGTCATCTCTCTGGGGTGTATCTGGT

Legend: **TCF sites** CDX sites CAG sites **SOX sites**

## B

| Probe | Sequence |
|---|---|
| WT | CAATTTAACAATGTGTTCTTTGTCCCACAA |
| 5’ mut | CAATTTAA**ACC**TGTGTTCTTTGTCCCACAA |
| 3’ mut | CAATTTAACAATGTGTTCT**GGT**TCCCACAA |
| 5’+3’ mut | CAATTTAA**ACC**TGTGTTCT**GGT**TCCCACAA |

**Fig S2. The c-Myc-335 WRE is bound and regulated by SOX9.**
**A)** Sequence of the c-Myc-335 WRE with regulatory motifs annotated. Positions shown are based on the GRCh38.p13 primary assembly of the *Homo sapiens* genome. **B)** Sequences (5’-3’) of the SOX site probes used for EMSA. The SOX sites in the WT and corresponding locations in the mutant probes have been underlined. Mutated bases are emboldened.

## A

Myc-8 enhancer sequence:
Chr8:127742878-127744069

GGTCTTCAGCTCCCTGTACTTTGGGATTTTAATCTACCACCACCCATAAATCAATAAATAATTACTTTCTTTGACTCTGACTCCTAGAA
TAATCTATTCAAAACCTTAATGTCTTTTTCTTGATCCTTCTTTTGAGTCCTAAGTACaCATTACAGCTTCAAATTGGCACGTCATATAG
GCGAA**TTTCAAAGG**G**AGATGCAA**TCCACAGAAGTATAGTA**GTTCAAAGG**GTTACAAA**AGCAAGGC**G**CTCTTAAA**CAGCTCAGTCTTTGC
C**CCTTTGTGG**CCTAGGGCTGGAGTGCAGCTCTGGGGTGACTCACTTGGGAATCGGGAAGGTGTTAGTCTGAATCACTAAGTCCAGGCAA
GCCCTCAGAATAGGAGAGAGTGTTCCTAGCAAGGAAAACAACTCTCCATTCCAAATAATCAGGAAAGAACTTTAGGGATGTGGAGCTTG
GCTATGGGAATAGAAAGGAACCATTCCAAGTGCCTATTAGGCCGCTCTTACCTTTACTGAGCCAGAGAATGGCTCTGAAAACAGGACAG
ATGCCAACTTCCTTCCCGAAAGTCAGGCTGATCTTGACCACAATACAAATTGGCCCTTAGAGCCTATACAGGGAGTCCCAGGGGTCTCT
GCCATTGTGCAACCTATTTTGTAGATAATAATCAAGAATCGGACGTGAAGGGGAGGAGTTTGCAACTTGGTCAGGAATGTATAAGAAGG
AATAAGCTAATTCTGACTATGCCCTTTATCCATGACACTATCCAGGAATTAATGACTCTCCCAGAGGATTCCTGGAATGATTTTGTTGA
GGGATGGAATGTATAAAGAGGAAGGAAGTGTTATTTTATGCTGCCATTTGGAAG**CAACAAAGG**AGATCAACAGTATGAAAACAATCAAT
CAAATTTGAAAATGAACAAAGTTTTCACAATCCCAGCCTAATACTTAGAGAGCTCACAGCTTGGATGCATAAGTAAAGAGTTCTCTGCT
GGTCTTTAAGACAAACTCTCACACAAAACTTGGGAAAAAGGACAAAAATGTTGCATTAGGGGGTTTTCTGTGGTTTGTTTGCAATAACT
ATAATTGGCTCA**ATCAATAA**T**TATTTTTT**AGTATACACACTAAGGGCCCCTGTAGCATTTTTTCCCATCG**ATAAATAA**T**CCTTAGTC**TA
GAAAATGCCGAGGGATGTTCTCCACCCTTGTCT

## B

Myc-29 enhancer sequence:
Chr8:127706818-127707901

AAAAGGTTGGGCTGGTTTCAAATGTTCTCAG**AACCAAAGC**AACACACGCCTTTCAGAGCGCTGGAGAAAATGCTGTTGAACAAGCTCTC
ATGGCTCAACAGTGGAGATCTGAACAATGTATCTGACTATTAACATAATTTGAGAAGAGAGGCTCCCTCTTAACCTGTATTTTTGGCTC
ACTCAAAAAAAGTGCTCCTGTCAGCCTTCAATTTCTCTACCTGGGTCTCTAGTCTGAGGACAGCATCCTTTTGGAAGACTTCAGCAATT
TCTGTGGAGCCATGTTTAAATTGGGCAAGTGGAAAACTAACTCCCTTTA**AGAAAAGT**G**GCATTGTT**TTGCAGTCAGGTAATTCCACGTG
GGCTCACTGGTTACTTTCCATGGGAATAG**TCTTTGAGGAAGAAAGG**G**CCTTTGTTG**AGATTTGTTTTGTTTTGAGAACAGTCAATTGGT
TCTTTCTACTCTTCCTTCTAGGGCGCCAATCTTTCCAAGGATAA**CCACAAAGT**GCAAGCAAAAATAATGAAAAAGGAAAGAAAGAAAGA
AAATCTGTATGCAAACCCAGGTTGTCATTATCTGGCGTTGTTTTCGGCCATTCTTGTCTTCTGACTCCTACCTGGCACACAGATCACTC
CTGCCCACTCCTGCCCATTACTCTCTATCTTGTACACGCCTTGCTGGTACATATGGAGGACCATGGAAGAAAAGCATTGCTGTAAATTT
CCTCCTT**CTCAAGGA**A**TCCTGCTC**CCTGAGTTTCTGCTTAAGTGACTTCTCACCTTCAAACTCACTGAGGTTATTCTTGTTTGGCTTCA
GATAAGTGGCTGAGATTTTAATTGTTCTCAATTGGTCTGTTTATCAGGGTAACTGTTTCAGTTTTCTTCAAGACAAATGATTAGATCCT
TCGGTCTTTATCAGTATATTAAGTATGTTTTTCAGGTGTGGACTTAACTTTGCCCTCCACTCCCAGCAACCGTTTCCTTTCATAACTTG
GGTGTAGTTTAAACACACACACACACACACACACACACACACACACACTTGCTCAGAGTTATTTTGGAAAGTGTATTCTTTACCAGTGA
CACCAACCTCAAAAGA

## C

Myc-521 enhancer sequence:
Chr8:127214627-127216810

ACAGTGTTAGACA**GTACAAAGG**GCATTTTTACCTCCAACACATCTTTCATGAAGAGTTTTTTATCTAGGTTTCATGGAGCCCTACGGTA
AATAAATTTGCCCAAGTTCTAACAAGATTCTGGAATCTCATATGTGGGTATGTTCATGTTGGGGTGTGTGTGTGTGTGTGTATGTGGGG
TGTATGGGTGTCTGTGTTTTTCCTCTGAAAAGCACCATAGTCTCTGCCACATTTTTAAAAAGGTGCATAAC**CCTTTAAAC**AGGAACCAT
TTACTTTAAATGGAATCAGAAATCTCACTATAAAAAAGCAATAATATTAATTTCTACTGTGTGTGTCTCTTGCCTTTTATTCCATGTTC
ACAATTCATCTTATTTTCATTTAAGACAACCAACTGGGCTAGTTAGAGCCGAGATTTGAGCCCAGTTTTTCTCACTGACTCTAAAACCT
ATCCATGCTCCTAAACCTCCTCACCATTGGAGGGCATTGCTGTTTACCCTTTCAGTTTCAGCTGTACTATCAAAAGCAGCAAATCAAGG
GCAGGGACCACAGCTTGACCTTAAACTATTCAACTGCA**CTTAATTA**C**CCATTGTA**GTGGAAGCACATAAATACAGGATGTCCCCTCACT
CTGCCATTCTGCTTTGCATCATTATCAAAA**AGACAAAGG**TAATTCATGTCTAATTACTCACTCACCGCAAGGTCAGAATATTTATTTTT
ATCATACTTGTGATTTTTAGGATTAGTGATATCAGTATTTATTCATGGCTAACCTAAATCAAACATCCTGTTGATATCTGTAAAGAGCC
ATATAGTTATTGTCGCAGGTATGCTGAGTCACAAAGCAAATACAAAGTTTATGAGTTGACAGGGAAAAGGCTGCAGAGAGGTAGAGATA
AGTGTGCCTGGGTAATCACAAGTGTCTTGTGAAATATACACATTGTTTATTTATATTGTCCATTTATCAAG**CTCAATGG**G**GGCTTGGG**A
AACTGAAAATAATCAAAATAGGAAAAATTATTGGTATTATTTTCTAAATTTGCTGCTGGTGCTGACTTTTTGGTTCAATTTATTAAATA
CTTACTTTAGAAATTATTCTCTGTTTTTAGTGGAATAGA**GCTTTGACC**AGCTTGAGAAAGACAAAAGGAGAAGTAGAAGAGAAGGGGAA
ATAAATGAGGCTCATGTGACTGGGATTAATTTACTCAGAGATCTGGAGTAGCAATAACTTGTGTCTTTCCAAAACCCCCGCAGTCAATT
AATGAGATCAATTTAAAGTGGCAGTATTGAAAGGGAAAACCCAAACACTTAGATATAGCAGTATGCAACTGACCTATTCCTTTTTCCAT
TCAGATTCAGTCAGTCAAATATATCTTATTG**ACTTTGACT**ACTGTCATCATAACTGTGTTAGCTAATTGT**CTGAAGTT**T**TCATTCAT**TC
TCTGTCTTTGCAGATGTCACTGCTATTCAATCTGTATTTCCTTGCTCTGTCTCGCTGTCTA*TCTGCACTAACAGGATATACACGTGTG
TGTACACATTTTTTTTTCTTAGTCTATCCCATTGTTTCTCTTGATGTCCTTGGACAGATCAGCAGTCTGAAAATCAGGCTTCTAATAGA
GAGTCTTACTAGAAGTAACTAGCTGTGAAACTTGGGCAAGTATTTGACCTCTCCCTTCCCCCATCTATGAAAGGAAGATGTGAAAACAA
ATGGTTCTGTGGGTCCTCTCCCACTTATTCACTCGATATTTCTATAATTTTCTATCTAGTTTCCATGGAGCACAAGGTCATTAGTAATG
TGCCCCATGACTCTCAGATTAATCAGACAAGCCAACACTGGGGGAGATGT**GAACAAAGT**CTTAGAGTCTAGTTGATAGTAAGAATATGA
AATGAATATGACTCACTGATGTGATATAGTCTTCAAAAAACTAAATTTGGTGTTTGAATGCGTAGAAATATATTAGGTTTTACAACATA
AAATTTCCATTTTGGTAGATGGAAAATGTTG**AATAAAGG**C**AATTTCAT**AAGGTTCATAGTCTGTAGAGTGAAATGGAAGTGGAAAAACC
TC**ACTTTGCTC**TGTTCTGCTCTACTTAGCTCATACTTGAAGTAATCATTCAGTTCTTGATGCCATACAGACAAAGAAGGTAGTGGACAA
AAAGAGACACACATATTCAGTAGATAGCAACCAAGCTGGGAGGGGAAC

**Fig S3. Sequences of *MYC* WREs bound and regulated by SOX9.**
**A)** Sequence of the Myc-8 WRE with regulatory motifs annotated. **B)** Sequence of the Myc-29 WRE. **C)** Sequence of the Myc-521 WRE. Positions shown are based on the GRCh38.p13 primary assembly of the *Homo sapiens* genome. Putative TCF and SOX9 binding sites are annotated.

**A**

*Defa5* promoter sequence:
Chr8:7057354-7056678

GATGCATTCAGATCACACCAACTCCTTGAACTAAATCCGAATTTTTATTTTAATCTGATAAACTTGGCCTA
CTATTTTACTGAACTCATTTCCCCTATAGCCTGATAAGGTCATTGACCTCTCCATACTGGCACCAGCGGGA
GACTACTCACCTCGAGATCTCAAAAGCCTCCTACATGAGGTTAGTAATATCCCTGAATCCTGCAATGAATT
AACTCTCTACTCCACTGGGTCCCAGGTCTGCCCCCAGAGAGTCATCCAGAGAGTACCAGGGACCATCTTCA
GAAAACAAGAGGCATTTGATCCCCAAACTTCTTGAATGAAAGCGCTGTTGTTTTTCTTTTTTGAATATATA
AAAGTAAATACTCAAGCAGATGGGAAACAGAACAGGATAGTAATACCCTTATCATCATTAACACCTTGGAT
CAAGAAGAGGCATTAAGCATACAGACTCACGCTTTGATGAAAGCTGGGAGAAAGAGGAGCATCAAAGGGAT
CTTGAGAACAAAGGCAGTCCTTCCCCTCCCAATCACATGCCCACCTCCTCTCACTGCAGCTTCTGTCTCAG
GTCTTCTCCCAGCAGAGCTATAAATCCAGGCTGACTCCTCACTCCCCAC**ATATCCACTCCTGCTCTCCCTC**
**CTGCAGGTGACCCCAGCCAGAGGACCATCGCCATCCT**

Legend: TCF sites CDX sites CAG sites SOX sites **Exon**

**B**

Defa6 promoter sequence:
Chr8:6926680–6926025

TCACATGGGTTTCTTGAAATAAATCTTTTGGCTTTAGTTTTACTAACTCTTTACCTAGTATCCCACTGAGT
TCTTTTCCCTTATAGGCTGATAAGGTCATTATCTTCTCCACACTGTGCCCCACAAGAGCCTATTCACCCCA
TAATCTCAAAGCCTCCTTCATGAGGGCAGAAATGCCCCCTGAATCCTGCAATGAATTAACTCTCTACTCTA
GCGGGATCCAGCTCTGGCCTCAAGGTCTAGACCTCCAGAGAGTGGCCAGCCCCACCTTCAGAAAATAAGAG
GCATTTGATTCCTGAAATTATTCATTGAAAGCACTGTTCTTTTCTTTTTTGAATATTAACAAGTAAATATT
CCAGCAGATGGAAAACAGGACAATGTAACACTGTTCTTATCATCACTATCAGCTGGGACCAGAACAGACAC
TCAATAAACAGCCTCACACTACAATGAAGCTTGGAGAACAAAGGAGCATCAAAGGGACATGGAGGGCAAGG
GTAGCTCTTCTGCTCCCCAATCACATGCACTCCCCGTCTCACCGCAACATCTGTCCCTGAGCCTTCTCCCA
GCAGACCTATAAATCCAGGCTGGCTCCTCACTCCCC**ACACATCTGCTCCTGCTCTCTCTCCTCCAGCGACC**
**CTAGCCAGtAGAACCCT**

**C**

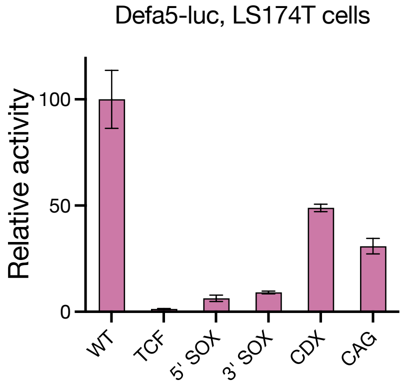


**D**

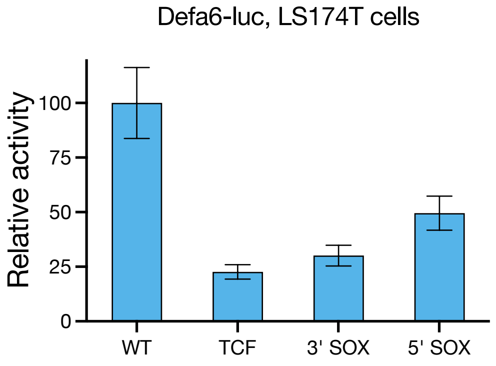


**Fig S4. Synergistic upregulation of Paneth cell defensins by Wnt signalling and SOX9.**
**A)** Sequence of the *Defa5* WRE with regulatory motifs annotated. The region between nucleotides -617 to +60 with respect to the transcription start site (TSS) were cloned to make the reporter. The ATG at +40 was changed to AGT in the reporter. **B)** Sequence of the *Defa6* WRE with regulatory motifs annotated. Positions shown are based on the GRCh38.p13 primary assembly of the *Homo sapiens* genome. The region between nucleotides -604 to +52 with respect to the TSS were cloned into the promoter. ATG at +41 was changed to AGT in the reporter. **C)** Luciferase assay data showing a reduction in Defa5-luc activity in LS174T cells upon the mutation of TCF, SOX, CDX, or CAG sites. **D)** Luciferase assay showing a reduction in Defa6-luc activity in LS174T cells upon mutation of TCF or SOX sites. Data in **(C)** and **(D)** are shown as mean ± s.d. of triplicates.

# A

### TOPflash

GGTACCTGAGCTCCACCGCGGTGGCGGCCGCTCTAGAACTAGTGGATCCCCCGG**GAGATCAAAGG**GGG
T**AAGATCAAAGG**GGGT**AAGATCAAAGG**GGTCGG**GAGATCAAAGG**GGGT**AAGATCAAAGG**GGGT**AAGAT**
**CAAAGG**GGTCGACCTCGAGGATATCAAGATCTGGCCTCGGCGGCCAAGCTT

# B

### SOXflash

AAATAGGCTGTCCCCAGTGCAAGTGCAGGTGCCAGAACATTTCTCTGGCCTAACTGGCCGGTACCTGA
GCTCGCTAGCCTCGAGATTT**AACAATGT**GT**TCTTTGTC**CCACATTT**AACAATGT**GT**TCTTTGTC**CCAC
ATTT**AACAATGT**GT**TCTTTGTC**CCACAGATCTGGCCTCGGCGGCCAAGCTT

# C

### TOP/SOX

ACGAAACAAAACAAACTAGCAAAATAGGCTGTCCCCAGTGCAAGTGCAGGTGCCAGAACATTTCTCTG
GCCTAACTGGCCGGTACCTGAGCTCGCTAGCCTCGAGTCGG**AACAATGT**G**TTCTTTGT**GTAC**GAGATC**
**AAAGG**GGGT**AACAATGT**G**TTCTTTGT**GTAC**AAGATCAAAGG**GGTCAAGCTT

Legend: **TCF sites** **SOX sites**

**Fig S5. Sequence determinants of synergistic activation by Wnt signalling and SOX9.**
Sequences of the **A)** TOPflash, **B)** SOXflash, and **C)** TOP/SOX synthetic reporters with TCF and SOX sites annotated.

**A**

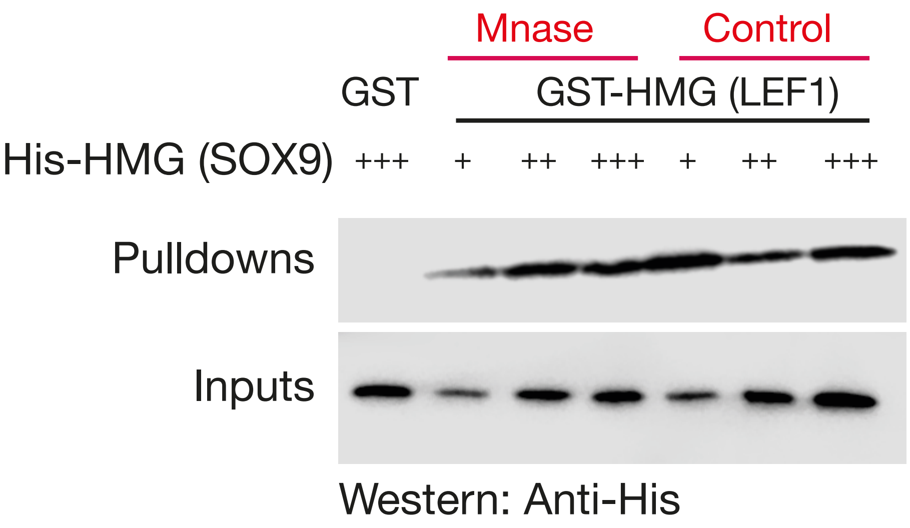


**B**

| Uniprot ID, species, protein | Sequence | Positions |
|---|---|---|
| Q04887<br>Mouse SOX9 | KLADQYPHLHNAELSKTLGKLWRLLNESEKRPFVEEAERLRVQHKKDHPDY | 122-<br>172 |
| P48436<br>Human SOX9 | KLADQYPHLHNAELSKTLGKLWRLLNESEKRPFVEEAERLRVQHKKDHPDY | 122-<br>172 |
| A0A337SCT0<br>Cat SOX9 | KLADQYPHLHNAELSKTLGKLWRLLNESEKRPFVEEAERLRVQHKKDHPDY | 122-<br>172 |
| P48434<br>Chick SOX9 | KLADQYPHLHNAELSKTLGKLWRLLNESEKRPFVEEAERLRVQHKKDHPDY | 122-<br>172 |
| Q6F2E7<br>X. tropicalis SOX9 | KLADQYPHLHNAELSKTLGKLWRLLNEGEKRPFVEEAERLRIQHKKDHPDY | 122-<br>172 |
| Q9DFH1<br>D. rerio SOX9b | KLADQYPHLHNAELSKTLGKLWRLLNEGEKRPFVEEAERLRVQHKKDHPDY | 108-<br>158 |
| Q9DFH2<br>D. rerio SOX9a | KLADQYPHLHNAELSKTLGKLWRLLNEVEKRPFVEEAERLRVQHKKDHPDY | 124-<br>174 |
| Q9VA17<br>D. melanogaster SOX100B | VMSKQYPHLQNSELSKSLGKLWKNLKDSDKKPFMEFAEKLRMTHKQEHPDY | 94-<br>144 |
| Q23045<br>C. elegans egl-13 | KILKAYPDMHNSNISKILGSRWKGMSNSEKQPYYEEQSRLSKLHMEQHPDY | 346-<br>396 |

**Fig S6. Highly conserved residues mediate a DNA-independent TCF/SOX9 interaction.**
**A)** Western blot analysis of a GST pulldown assay showing that the interaction of the HMG domain of LEF1 (GST-tagged) with the HMG domain of SOX9 (His tagged) is unaffected by treatment with Mnase, suggesting that it does not require DNA. **B)** Sequence comparison of SOX9 proteins across species showing conservation of residues mediating the TCF/SOX9 interaction.

| Protein | HMG sequence | Positions |
|---|---|---|
| SOX9 | RRKLADQYPHLHNAELSKTLGKLWRLLNESEKRPFVEEAERLRVQHKKDHPDYK | 120-173 |
| SOX8 | RRKLADQYPHLHNAELSKTLGKLWRLLSESEKRPFVEEAERLRVQHKKDHPDYK | 117-170 |
| SOX10 | RRKLADQYPHLHNAELSKTLGKLWRLLNESDKRPFIEEAERLRMQHKKDHPDYK | 119-172 |
| SOX7 | RKRLAVQNPDLHNAELSKMLGKSWKALTLSQKRPYVDEAERLRLQHMQDYPNYK | 60-113 |
| SOX17 | RKRLAQQNPDLHNAELSKMLGKSWKALTLAEKRPFVEEAERLRVQHMQDHPNYK | 83-136 |
| SOX18 | RKRLAQQNPDLHNAVLSKMLGKAWKELNAAEKRPFVEEAERLRVQHLRDHPNYK | 100-153 |
| SOX4 | RRKIMEQSPDMHNAEISKRLGKRWKLLKDSDKIPFIREAERLRLKHMADYPDYK | 74-127 |
| SOX11 | RRKIMEQSPDMHNAEISKRLGKRWKMLKDSEKIPFIREAERLRLKHMADYPDYK | 64-117 |
| SOX12 | RRKIMDQWPDMHNAEISKRLGRRWQLLQDSEKIPFVREAERLRLKHMADYPDYK | 55-108 |
| SOX5 | RRKILQAFPDMHNSNISKILGSRWKAMTNLEKQPYYEEQARLSKQHLEKYPDYK | 571-624 |
| SOX6 | RRKILQAFPDMHNSNISKILGSRWKSMSNQEKQPYYEEQARLSKIHLEKYPNYK | 636-689 |
| SOX13 | RRKILQAFPDMHNSSISKILGSRWKSMTNQEKQPYYEEQARLSRQHLEKYPDYK | 439-492 |
| SOX1 | RRKMAQENPKMHNSEISKRLGAEWKVMSEAEKRPFIDEAKRLRALHMKEHPDYK | 66-119 |
| SOX2 | RRKMAQENPKMHNSEISKRLGAEWKLLSETEKRPFIDEAKRLRALHMKEHPDYK | 56-109 |
| SOX3 | RRKMALENPKMHNSEISKRLGADWKLLTDAEKRPFIDEAKRLRAVHMKEYPDYK | 154-207 |
| SOX14 | RRKMAQENPKMHNSEISKRLGAEWKLLSEAEKRPYIDEAKRLRAQHMKEHPDYK | 23-76 |
| SOX21 | RRKMAQENPKMHNSEISKRLGAEWKLLTESEKRPFIDEAKRLRAMHMKEHPDYK | 23-76 |
| SOX15 | RRQMAQQNPKMHNSEISKRLGAQWKLLDEDEKRPFVEEAKRLRARHLRDYPDYK | 64-117 |
| SRY | RRKMALENPRMRNSEISKQLGYQWKMLTEAEKWPFFQEAQKLQAMHREKYPNYK | 75-128 |
| SOX30 | RPALAKANPAANNAEISVQLGLEWNKLSEEQKKPYYDEAQKIKEKHREEFPGWV | 352-405 |

**Fig S7. Conservation of TCF-interacting residues across human SOX9 proteins.**
When mutated in SOX9, D125,168,171 prevent it from binding to the HMG domain of TCFs. Amino acid residues in other human SOX family members that have acidic (D/E) residues at the corresponding positions are highlighted.

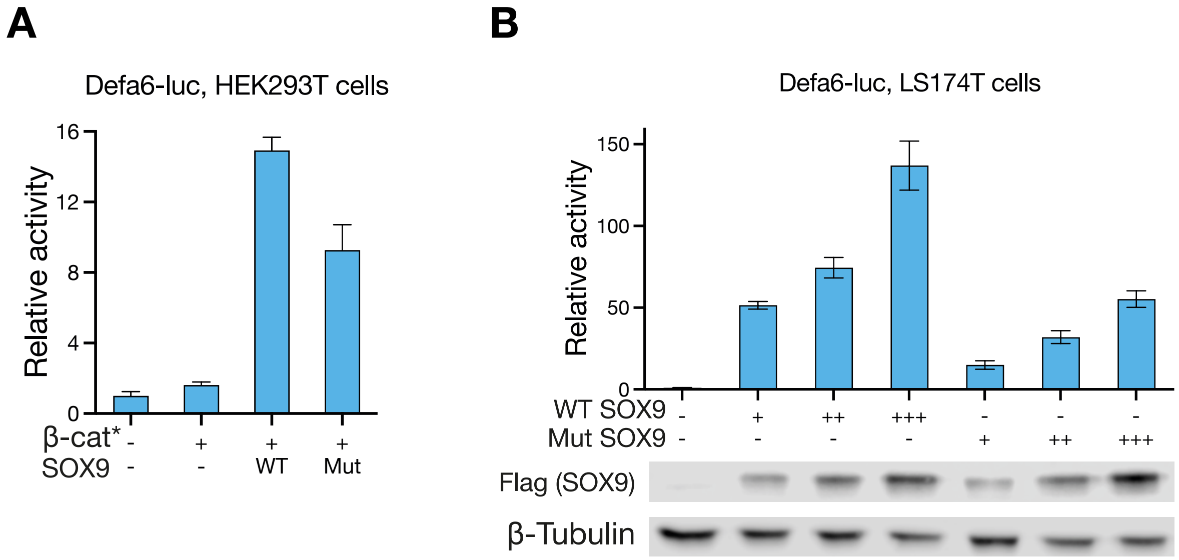


**Fig S8. Non-DNA contacting TCF-interacting residues play a role in the activation of Wnt-SOX9 activated enhancers.**

**A,B)** Luciferase assay data showing the activation of Defa6-luc by β-catenin* and WT or Mut SOX9 in HEK293T cells **(A)** and by WT or Mut SOX9 in LS174T cells **(B)**. Western blots in **(B)** show relative expression levels of WT and Mut SOX9 with Tubulin as loading control. **(C)** When mutated in SOX9, D125,168,171 prevent it from binding to the HMG domain of TCFs. Amino acid residues in other human SOX family members that have acidic (D/E) residues at the corresponding positions are highlighted.

| Construct | Target gene | pGIPZ clone ID | Target sequence |
|---|---|---|---|
| Scr | Scrambled negative control | RHS4346 | ATCTCGCTTGGGCGAGAGTAAG |
| SOX9-1 | SOX9 | V2LHS_92504 | TAAGACTGCAGTGAACAAG |
| SOX9-2 | SOX9 | V2LHS_11387 | ATGTCTTGAAGGTTAACTG |

**Table S1. Targeting sequences and clone IDs of the pGPIZ RNAi clones used to create LS174T cell lines expressing scrambled (Scr) or SOX9-targeting shRNAs.**
"Target sequence" denotes the sequence of the mature antisense siRNA.

| Construct | Site | Original site | Mutated site |
|---|---|---|---|
| 5' SOX | SOX9 | AACAATGT | AAaccTGT |
| 3' SOX | SOX9 | TCTTTGTC | gaTgTGTC |
| TCF+SOX | SOX9 (3' SOX) | TCTTTGTC | gaTgTGTC |
| | TCF | CCTTTGAGC | CCTgTtcGC |
| | | GATGAAAGG | GAgtAcAGG |
| | | GTACAAAGA | GTACAcAtc |
| | | AATCAAAGG | AAgaAcAGG |

**Table S2. Site-directed mutagenesis of the c-Myc-335 reporter**

| Reporter | Site | Original | Mutated |
|---|---|---|---|
| **Defa5** | **TCF** | GCTTTGATG | GCTGTGCGG |
| | | CATCAAAGG | CCGCACAGG |
| | | GAACAAAGG | GCCCACAGG |
| | **5' SOX** | ATGAAAGC | ATTCCAGC |
| | **3' SOX** | CTGTTGTT | CTGCGTTT |
| | **CAG** | TACTGA | TTCACA |
| | | TACTGG | TTCACG |
| | | CACTGG | CTCACG |
| | | ACAGAC | AGACTC |
| | | GGCTGA | GCCACA |
| | | CCAGCC | CGTGGC |
| **Defa6** | **TCF** | TCTCAAAGC | TCGTACAGC |
| | | ATTGAAAGC | AGGGACAGC |
| | | GAACAAAGG | GCCCACAGG |
| | | CATCAAAGG | CCGCACAGG |
| | **5' SOX** | CTATTCAC | CGATGCCC |
| | | TCCTTCAT | TACTGCCT |
| | | GCTTGGAG | GATTTGCG |
| | | ACATGGAG | AAATTGCG |
| | **3' SOX** | CACAAGAG | CAACCGAG |
| | | CTCAAAGC | CTACCAGC |
| | | TACAATGA | TAACCTGA |

**Table S3. Site-directed mutagenesis of the *Defa5/6* reporters**

| Target | Primer |
|---|---|
| Defa5 | ACCTCAGGTTCTCAGGCAAG |
| | CTGATTTCACACACCCCGGA |
| G6PD (control) | GAGGCCGTGTACACCAAGAT |
| | TCAGGGAGCTTCACGTTCTT |
| Myc | TACAACACCCGAGCAAGGAC |
| | AGCTAACGTTGAGGGGCATC |

**Table S4. Primers used for RT-qPCR**

| Target | Primer |
|---|---|
| c-Myc-335 (Peak) | TCCTATCTCAGCTCCCTATCCA |
| | CTGCTGAGCTCAAAGGACGA |
| c-Myc-335 flank 1 | GGCTCTCACCCTTCAACCAA |
| | GCTTGTGACTTAGCCTGGGT |
| c-Myc-335 flank 2 | GGCTCTGGTTGGGGGTTTTA |
| | TGAGGCGGAAGTCAACACAG |
| Control | CGATTCAGTGCCGCATTAGC |
| | TAATCGGCTGGATCTCCCCA |

**Table S5. Primers used for ChIP-qPCR**
